# A genome-wide screen in macrophages identifies new regulators of IFNγ-inducible MHCII that contribute to T cell activation

**DOI:** 10.1101/2020.08.12.248252

**Authors:** Michael C. Kiritsy, Laurisa M. Ankley, Justin D. Trombley, Gabrielle P. Huizinga, Audrey E. Lord, Pontus Orning, Roland Elling, Katherine A. Fitzgerald, Andrew J. Olive

## Abstract

Cytokine-mediated activation of host immunity is central to the control of pathogens. A key cytokine in protective immunity is interferon-gamma (IFNγ), which is a potent activator of antimicrobial and immunomodulatory effectors within the host. A major role of IFNγ is to induce major histocompatibility complex class II molecules (MHCII) on the surface of cells, which is required for CD4^+^ T cell activation. Despite its central role in host immunity, the complex and dynamic regulation of IFNγ-induced MHCII is not well understood. Here, we integrated functional genomics and transcriptomics to comprehensively define the genetic control of IFNγ-mediated MHCII surface expression in macrophages. Using a genome-wide CRISPR-Cas9 library we identified genes that control MHCII surface expression, many of which have yet to be associated with MHCII. Mechanistic studies uncovered two parallel pathways of IFNγ-mediated MHCII control that require the multifunctional glycogen synthase kinase 3 beta (GSK3β) or the mediator complex subunit MED16. Both pathways are necessary for IFNγ-mediated induction of the MHCII transactivator CIITA, MHCII expression, and CD4^+^ T cell activation. Using transcriptomic analysis, we defined the regulons controlled by GSK3β and MED16 in the presence and absence of IFNγ and identified unique networks of the IFNγ-mediated transcriptional landscape that are controlled by each gene. Our analysis suggests GSK3β and MED16 control distinct aspects of the IFNγ-response and are critical for macrophages to respond appropriately to IFNγ. Our results define previously unappreciated regulation of MHCII expression that is required to control CD4^+^ T cell responses by macrophages. These discoveries will aid in our basic understanding of macrophage-mediated immunity and will shed light on mechanisms of failed adaptive responses pervasive in infectious disease, autoimmunity, and cancer.

## Introduction

Activation of the host response to infection requires the coordinated interaction between antigen presenting cells (APCs) and T cells (1–3). For CD4^+^ T cells, the binding of the T cell receptor (TCR) to the peptide-loaded major histocompatibility complex class II (MHCII) on the surface of APCs is necessary for both CD4^+^ T cell activation and their continued effector function in peripheral tissues (3–5). Dysregulation of MHCII control leads to a variety of conditions including the development autoimmunity and increased susceptibility to pathogens and cancers (6–10). While MHCII is constitutively expressed on dendritic cells and B cells, the production of the cytokine IFNγ promotes MHCII expression broadly in other cellular populations including macrophages (11–14). The induction of MHCII in these tissues activates a feedforward loop wherein IFNγ-producing CD4^+^ T cells induce myeloid MHCII expression, which in turn amplifies CD4^+^ T cell responses (14–16). Thus, IFNγ-mediated MHCII expression is essential for protective immunity.

The IFNγ-dependent control of MHCII is complex (5, 12, 17–19). Binding of IFNγ to its receptor induces cytoskeletal and membrane rearrangement that results in the activation of Janus kinases 1 and 2 (JAK1 and JAK2) and STAT1-dependent transcription (20, 21). STAT1 induces IRF1, which then drives the expression of the MHCII master regulator, CIITA (22, 23). The activation of CIITA opens the chromatin environment surrounding the MHCII locus and recruits transcription factors, including CREB1 and RFX5 (5, 24). MHCII is also regulated post-translationally to control the trafficking, peptide loading, and stability of MHCII on the surface of cells (25–27). While recent evidence points to additional regulatory mechanisms of IFNγ-mediated MHCII expression, including the response to oxidative stress, these have not been investigated directly in macrophages (17).

In non-inflammatory conditions, macrophages express low levels of MHCII that is uniquely dependent on NFAT5 (15). While basal MHCII expression on macrophages plays a role in graft rejection, it is insufficient to control intracellular bacterial pathogens, which require IFNγ-activation to propagate protective CD4^+^ T cell responses (28–30). Many pathogens including *Mycobacterium tuberculosis* and *Chlamydia trachomatis* inhibit IFNγ-mediated MHCII induction to evade CD4^+^ T cell-mediated control and drive pathogen persistence (31–33). Overcoming these pathogen immune evasion tactics is essential to develop new treatments or immunization strategies that provide long-term protection (28). Without a full understanding of the global mechanisms controlling IFNγ-mediated MHCII regulation in macrophages, it has proven difficult to dissect the mechanisms related to MHCII expression that cause disease or lead to infection susceptibility.

Here we globally defined the regulatory networks that control IFNγ-mediated MHCII surface expression on macrophages. Using CRISPR-Cas9 to perform a forward genetic screen, we identified the major components of the IFNγ-regulatory pathway in addition to many genes with no previously known role in MHCII regulation. Follow-up studies identified two critical regulators of IFNγ-dependent CIITA expression in macrophages, MED16 and GSK3β. Loss of either MED16 or GSK3β resulted in significantly reduced MHCII expression on macrophages, unique changes in the IFNγ-transcriptional landscape, and prevented the effective activation of CD4^+^ T cells. These results show that IFNγ-mediated MHCII expression in macrophages is finely tuned through parallel regulatory networks that interact to drive efficient CD4^+^ T cell responses.

## Results

### Optimization of CRISPR-Cas9 editing in macrophages to identify regulators of IFNγ-inducible MHCII

To better understand the regulation of IFNγ-inducible MHCII we optimized gene-editing in immortalized bone marrow-derived macrophages (iBMDMs) from C57BL6/J mice. iBMDMs were transduced with Cas9-expessing lentivirus and Cas9-mediated editing was evaluated by targeting the surface protein CD11b with two distinct single guide RNAs (sgRNA). When we compared CD11b surface expression to a non-targeting control (NTC) sgRNA by flow cytometry, we observed less than 50% of cells targeted with either of the CD11b sgRNA were successfully edited (Figure S1A). We hypothesized that the polyclonal Cas9-iBMDM cells variably expressed Cas9 leading to inefficient editing. To address this, we isolated a clonal population of Cas9-iBMDMs using limiting dilution plating. Using the same CD11b sgRNAs in a clonal population (clone L3) we found 85-99% of cells were deficient in CD11b expression by flow cytometry compared to NTC (Figure S1B). Successful editing was verified by genotyping the CD11b locus for indels at the sgRNA targeting sequence using Tracking of Indels by Decomposition (TIDE) analysis (34). Therefore, clone L3 Cas9^+^ iBMDMs proved to be a robust tool for gene editing in murine macrophages.

To test the suitability of these cells to dissect IFNγ-mediated MHCII induction we next targeted Rfx5, a known regulator of MHCII expression, with two independent sgRNAs (35). We stimulated Rfx5 targeted and NTC cells with IFNγ for 18 hours and quantified the surface expression of MHCII by flow cytometry (Fig 1A and 1B). In cells expressing the non-targeting sgRNA, IFNγ stimulation resulted in a 20-fold increase in MHCII. In contrast, cells transduced with either of two independent sgRNAs targeting Rfx5 failed to induce the surface expression of MHCII following IFNγ stimulation. Thus, L3 cells are responsive to IFNγ and can be effectively used to interrogate IFNγ-mediated MHCII expression in macrophages.

**Figure 1.**
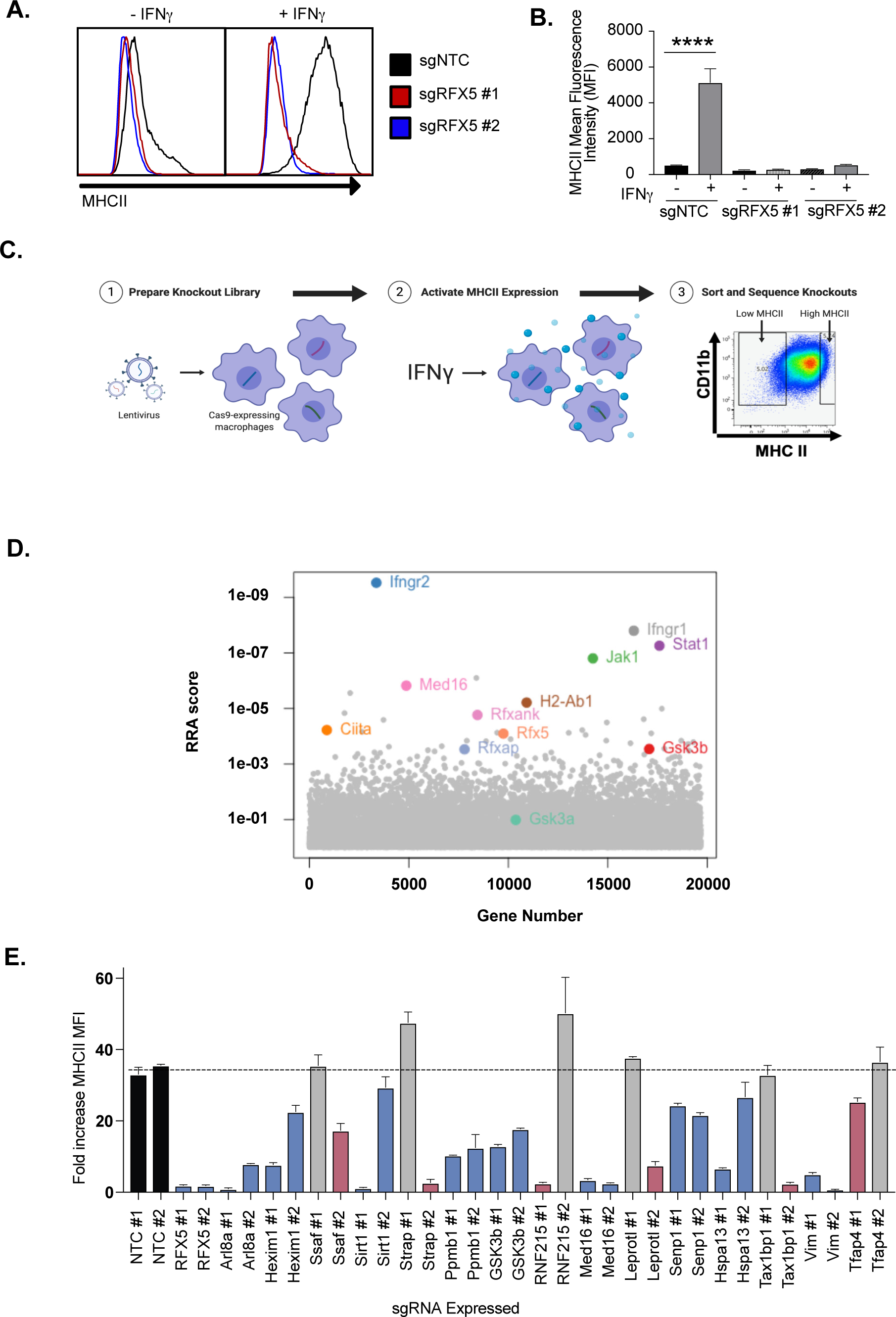
Genome-wide CRISPR Cas9 Screen Identifies regulators of IFNγ-dependent MHCII expression. **(A)** Cas9+ iBMDMs (Clone L3) expressing the indicated sgRNAs were left untreated or treated with IFNγ (6.25ng/ml) for 24 hours. Surface MHCII was quantified by flow cytometry. Shown is a representative histogram of MHCII surface staining and **(B)** the quantification of the mean fluorescence intensity (MFI) in the presence and absence of IFNγ stimulation from 3 biological replicates. **** p<.0001 by one-way ANOVA with tukey correction for multiple hypotheses. These data are representative of three independent experiments. **(C)** A schematic representation of the CRISPR-Cas9 screen conducted to identify regulators of IFNγ-inducible MHCII surface expression on macrophages. A genome-wide CRISPR Cas9 library was generated in L3 cells using sgRNAs from the Brie library (4 sgRNAs per gene). The library was treated with IFNγ and MHCII^hi^ and MHCII^low^ populations were isolated by FACS. The representation of sgRNAs in each population in addition to input library were sequenced. **(D)** Shown is score for each gene in the CRISPR-Cas9 library that passed filtering metrics as determined by the alpha-robust rank algorithm (α-RRA) in MAGeCK from two independent screen replicates. **(E)** The L3 clone was transduced with the indicated sgRNAs for candidates (2 per candidate gene) in the top 100 candidates from the CRISPR-Cas9 screen. All cells were left untreated or treated with 10ng/ul of IFNγ for 24 hours then were analyzed by flow cytometry. The fold-increase in MFI was calculated for triplicate samples for each cell line (MFI IFNγ+/MFI IFNγ-). The results are representative of at least two independent experiments. Candidates that were significant for two sgRNAs (Red) or one sgRNA (Blue) by one-way ANOVA compared to the mean of NTC1 and NTC2 using Dunnets multiple comparison test. Non-significant results are shown in Grey bars.

### Forward genetic screen identifies known and novel regulators of MHCII surface expression in macrophages

To define the genetic networks required for IFNγ-mediated MHCII expression, we made a genome-wide library of mutant macrophages with sgRNAs from the Brie library to generate null alleles in all protein-coding genes (36). After verifying coverage and minimal skew in the initial library, we conducted a forward genetic screen to identify regulators of IFNγ-dependent MHCII expression (Figure 1C and Table S1). The loss-of-function library was stimulated with IFNγ and 24 hours later, we selected MHCII^high^ and MHCII^low^ expressing cells by fluorescence activated cells sorting (FACS). Following genomic DNA extraction, sgRNA abundances for each sorted bin were determined by deep sequencing.

As our knockout library relied on the formation of Cas9-induced indels and was exclusive to protein-coding genes, we focused our analysis on genes expressed in macrophages under the conditions of interest, which we determined empirically in the isogenic cell line by RNA-seq (Table S2). We assumed that sgRNAs targeting non-transcribed genes are neutral in their effect on IFNγ-induced MHCII expression, which afforded us ∼32,000 internal negative control sgRNAs (37). To test for statistical enrichment of sgRNAs and genes, we used the modified robust rank algorithm (α-RRA) employed by Model-based Analysis of Genome-wide CRISPR/Cas9 Knockout (MAGeCK), which first ranks sgRNAs by effect and then filters low ranking sgRNAs to improve gene significance testing (38). We tuned the sgRNA threshold parameter to optimize the number of significant hits without compromising the calculated q-values of known positive controls that are expected to be required for IFNγ-mediated MHCII expression. Further, by removing irrelevant sgRNAs that targeted genes not transcribed in our conditions, we removed potential false positives and improved the positive predictive value of the screen (Figure S2A and S2B).

Guide-level analysis confirmed the ability to detect positive control sgRNAs which had robust enrichment in the MHCII^low^ population (Fig S2C). Using the previously determined parameters, we tested for significantly enriched genes that regulated MHCII surface levels. As expected, sgRNAs targeting known components of the IFNγ-receptor signal transduction pathway, such as Ifngr1, Ifngr2, Jak1 and Stat1, as well as regulators and components of IFNγ-mediated MHCII expression, such as Ciita, Rfx5, and Rfxank were all significantly enriched (Figure 1D) (5, 22). These results validated our approach to identify functional regulators of IFNγ-mediated MHCII expression.

Stringent analysis revealed a significant enrichment of genes with no known involvement in interferon responses and antigen presentation. To identify functional pathways that are associated with these genes, we performed KEGG pathway analysis on the positive regulators of IFNγ-induced MHCII that met the FDR cutoff (FigS2D) (39–41). However, gene membership for the ten most enriched KEGG pathways was largely dominated by known regulators of IFNγ signaling. To circumvent this redundancy and identify novel pathways enriched from our candidate gene list, the gene list was truncated to remove the 11 known IFNγ signaling regulators. Upon reanalysis, several novel pathways emerged, including mTOR signaling (Figure S2E). Thus, our genetic screen uncovered previously undescribed pathways that are critical to control IFNγ-mediated MHCII surface expression in macrophages.

The results of the genome-wide CRISPR screen highlight the sensitivity and specificity of our approach and analysis pipeline. To gain new insights into IFNγ-mediated MHCII regulation, we next validated a subset of candidates that were not previously associated with the IFNγ-signaling pathway. Using two independent sgRNAs for each of 15 candidate genes, we generated loss-of-function macrophages in the L3 clone (Figure 1E and S2F). MHCII surface expression was quantified by flow cytometry for each cell line in the presence and absence of IFNγ activation. For all 15 candidates, we observed deficient MHCII induction following IFNγ stimulation with at least one sgRNA. For 9 of15 candidate genes, we observed a significant reduction in MHCII surface expression with both gene-specific sgRNAs These results show that our screen not only identified known regulators of IFNγ-mediated MHCII induction, but also uncovered new regulatory networks required for MHCII expression on macrophages.

We were interested in better understanding the IFNγ-mediated transcriptional activation of MHCII to determine if a subset of candidates reveal new regulatory mechanisms of MHCII-expression. Based on the screen and validation results, we examined the known functions of the candidates that were confirmed with two sgRNAs, and identified MED16 and Glycogen synthase kinase 3b (GSK3β) for follow-up study. MED16 is a subunit of the mediator complex that regulates transcription initiation while GSK3β is a multifunctional kinase that controls signaling pathways known to regulate transcription (42, 43). Thus, we hypothesized that MED16 and GSK3β would be required for effective IFNγ-mediated transcriptional control of MHCII.

### MED16 is uniquely required for IFNγ-mediated CIITA expression

We first examined the role of MED16 in controlling IFNγ-mediated MHCII expression. MED16 was the sixth ranked candidate from our screen results, with robust enrichment of all 4 sgRNAs in the MHCII^low^ population (Fig2A). Our validation results confirmed that MED16 was indeed an essential positive regulator of MHCII expression (Fig1E). As part of the mediator complex, MED16 bridges the transcription factor binding and the chromatin remodeling that are required for transcriptional activation (44). These changes then recruit and activate RNA polymerase II to initiate transcription. While the core mediator complex function is required for many RNA polymerase II dependent transcripts, distinct sub-units of the mediator complex can also play unique roles in gene regulation (42, 44). To examine if MED16 was uniquely required for IFNγ-dependent MHCII expression, we probed our genetic screen data for all mediator complex subunits. None of the other 27 mediator complex subunits in our library showed any significant changes in MHCII expression (Figure 2B). To test the specific requirement of MED16, we generated knockout macrophages in Med16 (Med16 KO) and using two independent sgRNAs targeted three additional mediator complex subunits, Med1, Med12 and Med17 (Figure S3 and materials and methods). We treated with IFNγ and quantified the surface levels of MHCII by flow cytometry. In support of the screen results, Med1, Med12 and Med17 showed similar MHCII upregulation compared to NTC cells, while Med16 targeted cells demonstrated defects in MHCII surface expression (Figure 2C and Figure 2D). These results suggest that there is specificity to the requirement for MED16-dependent control of IFNγ-induced CIITA that is unique among the mediator complex subunits.

**Figure 2.**
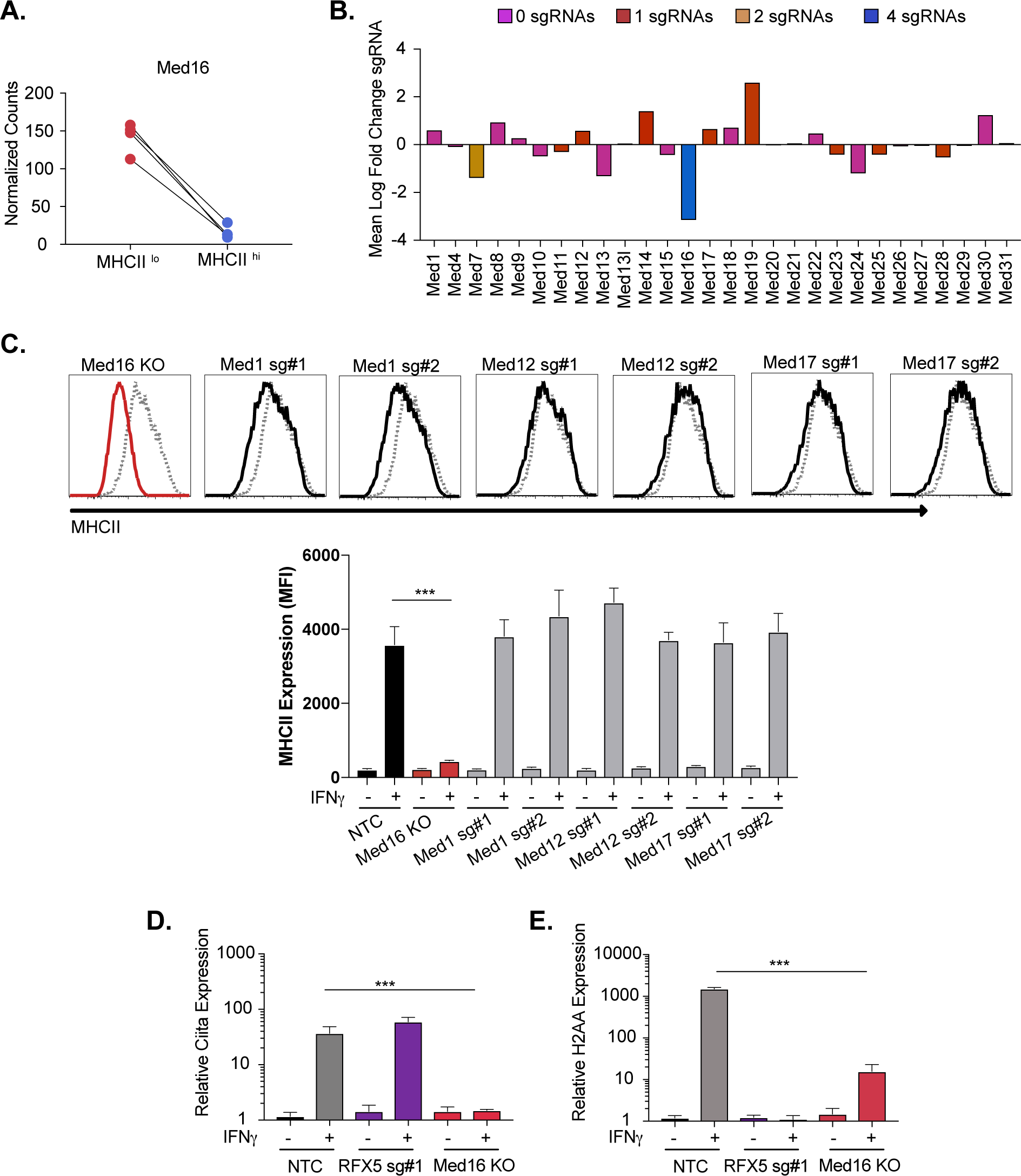
The mediator complex sub-unit MED16 is uniquely required for IFNγ-mediated MHCII surface expression. **(A)** Shown is the normalized mean read counts from FACS sorted MHCII^low^ and MHCII^hi^ populations for the four sgRNAs targeting MED16 within the genome-wide CRISPR-Cas9 library. **(B)** The mean of the log fold change (normalized counts in MHCII^hi^/normalized counts in MHCII^low^) for each mediator complex subunit that passed quality control metrics described in materials and methods. The bar colors indicate the number of sgRNAs out of four possible that pass the alpha cutoff using the MAGeCK analysis pipeline as described in material and methods. **(C)** Med16 KO cells or L3 cells targeted with the indicated sgRNA were left untreated or were treated with 6.25 ng/ml of IFNγ for 18 hours. Cells were then analyzed for surface MHCII expression by flow cytometry. Shown are representative comparing the MHCII surface expression of indicated mediator complex subunit (Black solid line) treated with IFNγ overlayed with NTC (Grey dashed line) treated with IFNγ. **(D)** Quantification of the MFI of surface MHCII from the experiment in (C) from three biological replicates. These results are representative of two independent experiments. **(E)** NTC L3 cells, RFX5 sg#1 cells, and Med16 KO cells were left untreated or were treated with 6.25 ng/ml of IFNγ. 18 hours later cells RNA was isolated and qRT-PCR was used to determine the relative expression of CIITA and **(H)** H2aa compared to GAPDH controls from three biological replicates. The results are representative of three independent experiments. ***p<.001 as determined one-way ANOVA compared to NTC cells with a dunnets test.

To understand the mechanisms of how MED16 regulates MHCII-induction, we assessed the transcriptional induction of MHCII in Med16 KO cells. In macrophages, the IFNγ-mediated transcriptional induction of MHCII subunits requires the transcriptional activation of CIITA that then, in complex with other factors like RFX5, initiates transcription at the MHCII locus (14, 17). To determine whether MED16 controls the transcriptional induction of MHCII, we stimulated NTC, Med16 KO and Rfx5 targeted cells with IFNγ for 18 hours and isolated RNA. Using qRT-PCR, we observed that loss of RFX5 did not impact the induction of CIITA, but had a profound defect in the expression of H2-Aa compared to NTC cells (Fig2E). Loss of MED16 significantly inhibited the induction of both CIITA and H2-Aa. These data suggest that MED16 controls the induction of MHCII through upstream regulation of CIITA.

### Loss of GSK3β prevents the IFNγ-dependent induction of CIITA

We next examined the mechanisms of GSK3β control of IFNγ-mediated MHCII expression in more detail. GSK3β is involved in many cellular pathways including mTor and Wnt (43, 45, 46). While GSK3β was previously suggested to repress collagen production via CIITA, no role in regulating IFNγ-mediated MHCII expression has previously been described (47). In addition to its high ranking in the screen and the strong effects of multiple sgRNAs (Figure 3A), pathway analysis uncovered a significant enrichment of genes within the mTor pathway, which controls GSK3β function, suggesting this signaling network is critical for MHCII expression (Figure S2E) (46). Our validation studies further showed that GSK3β is required for the effective induction of IFNγ-dependent MHCII (Figure 1E). To begin to understand the mechanisms controlling GSK3β-dependent regulation of MHCII expression we generated Gsk3β knockout cells (Gsk3β KO) and verified that the loss of Gsk3β strongly inhibited IFNγ-mediated MHCII surface expression (Figure 3B and Figure S3B).

**Figure 3.**
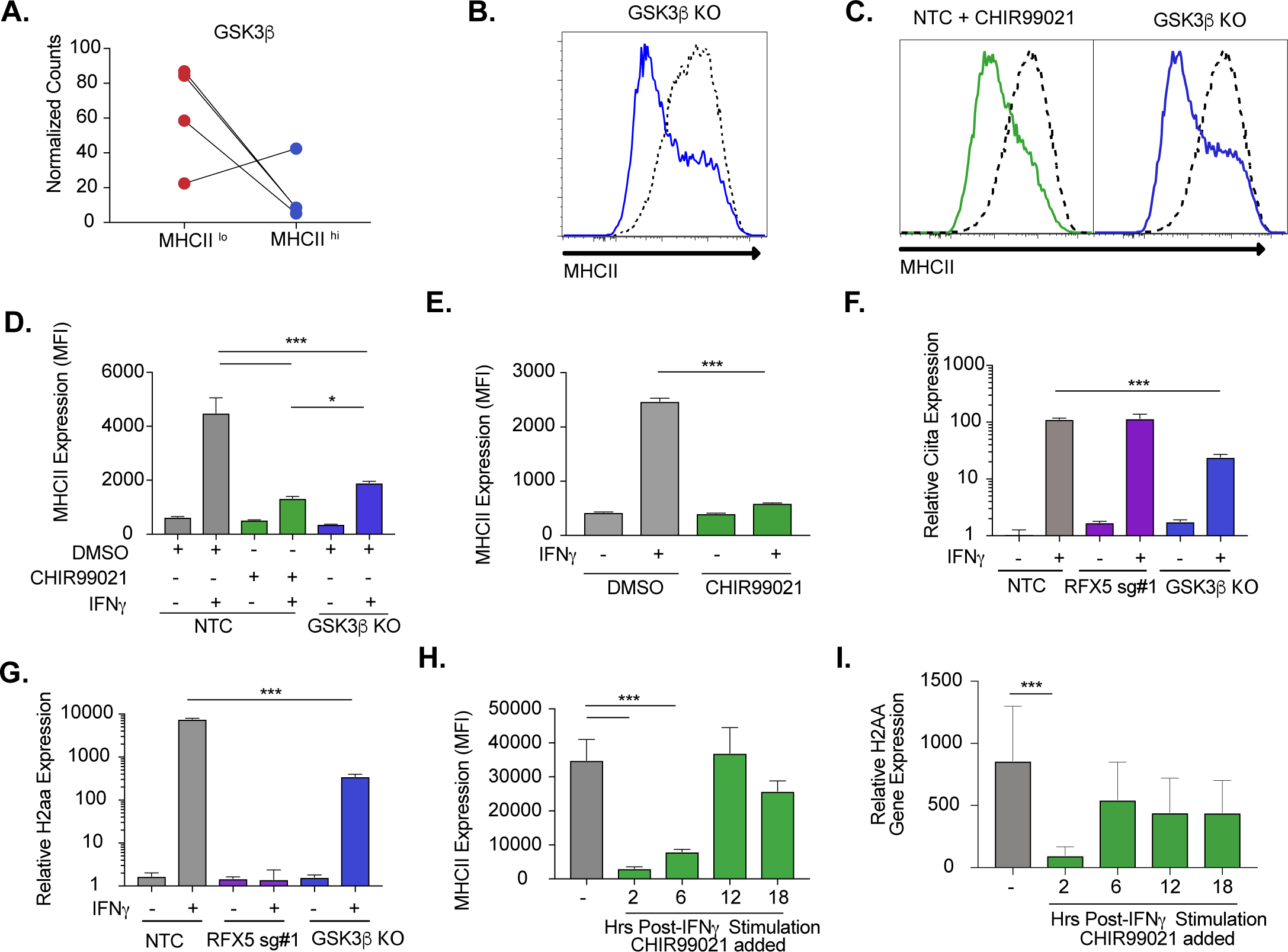
Inhibition of GSK3β results in decreased IFNγ-mediated CIITA and MHCII expression. **(A)** Shown is the normalized mean read counts from FACS sorted MHCII^lo^ and MHCII^hi^ populations for the four sgRNAs targeting Gsk3β within the genome-wide CRISPR-Cas9 library. **(B)** NTC L3 cells and Gsk3β KO cells were treated with 6.25 ng/ml of IFNγ. 18 hours later cells were stained for surface MHCII and analyzed by flow cytometry. Shown is a representative flow cytometry plot overlaying Gsk3β KO (blue line) with NTC (grey line). The results are representative of 5 independent experiments. **(C)** NTC L3 cells or Gsk3β KO were treated with DMSO or 10µM CHIR99021 as indicated then left untreated or stimulated with IFNγ for 18 hours. MHCII surface expression was then quantified by flow cytometry. A representative flow cytometry plot of DMSO treated IFNγ treated NTC cells is overlaid with either NTC CHIR99021 treated cells **(Left)** or Gsk3β KO DMSO treated cells **(Right)**. **(D)** The mean fluorescence intensity was quantified from three biological replicates. These results are representative of three independent experiments. **(E)** Bone marrow derived macrophages from conditionally immortalized HoxB8 progenitor cells from C57BL6/J mice were treated with DMSO or 10 µM CHIR99021 and left untreated or stimulated with IFNγ for 18 hours. The MHCII surface levels were then quantified by flow cytometry. Shown is the mean fluorescence intensity from 3 biological replicates in each condition. **(F)** NTC L3 cells, Rfx5 sg#1 cells, and Gsk3β KO cells were left untreated or were treated with 6.25ng/ml of IFNγ. 18 hours later cells RNA was isolated and qRT-PCR was used to determine the relative expression of CIITA and **(G)** H2-Aa compared to GAPDH controls from three biological replicates. The results are representative of three independent experiments. **(H)** Immortalized bone marrow macrophages were treated with IFNγ. Control cells were treated with DMSO and for the remaining cells CHIR999021 was added at the indicated times following IFNγ treatment. 24 hours after IFNγ stimulation the levels of surface MHCII were quantified by flow cytometry. Shown is the MFI for biological triplicate samples. **(I)** In parallel to **(H)**, 24 hours after IFNγ stimulation RNA was isolated and the relative expression of H2-Aa was quantified relative to GAPDH from biological triplicate samples. The data are representative of three independent experiments. ***p<.001 **p<.01 *p<.05 by one-way ANOVA with a Tukey Correction test.

To confirm the genetic evidence using an orthogonal method, GSK3β function was inhibited chemically using the well-characterized small molecule CHIR99021 (48, 49). NTC macrophages treated with either CHIR99021 or DMSO were stimulated cells with IFNγ and MHCII expression was analyzed by flow cytometry. Consistent with the genetic experiments, inhibition of GSK3β activity reduced the induction of surface MHCII, and was similar to a genetic loss of Gsk3β alone (Figure 3C and 3D). The GSK3β chemical inhibitor facilitated additional experiments in other cells that were not possible with Med16 KO cells. Thus, we repeated this experiment in primary bone marrow-derived macrophages from HoxB8 conditionally immortalized progenitor cells and observed identical results (Figure 3E) (50). Therefore, GSK3β activity is required for the effective induction of IFNγ-mediated MHCII in immortalized and primary murine macrophages.

We next examined if the IFNγ-mediated transcriptional induction of CIITA or H2-Aa were reduced in Gsk3β KO cells. Loss of GSK3β significantly inhibited the expression of both CIITA and H2-Aa after IFNγ-treatment compared to NTC controls. These data suggest that GSK3β, similar to MED16, is an upstream regulator of IFNγ-mediated MHCII induction that controls the expression of CIITA following IFNγ-activation.

These qRT-PCR studies (Figure 3F and 3G) suggested that GSK3β was required for the transcriptional activation of CIITA. We hypothesized that GSK3β inhibition with CHIR99021 would block MHCII expression only if the inhibitor was present shortly after IFNγ stimulation. To test this hypothesis, iBMDMs were stimulated with IFNγ then treated with DMSO for the length of the experiment or with CHIR99021, 2, 6, 12, and 18 hours post-stimulation. When MHCII was quantified by flow cytometry we saw a reduction in MHCII expression when CHIR99021 was added two or six hours after IFNγ (Figure 3H). CHIR99021 addition at later time points resulted in similar MHCII expression compared to DMSO treated cells. When the expression of H2-Aa mRNA was quantified from a parallel experiment, a significant reduction in mRNA expression was only observed in macrophages that were treated with CHIR99021 two hours following IFNγ-activation (Figure 3I). Thus, GSK3β activity is required early after IFNγ stimulation to activate the transcription of MHCII.

We were interested in understanding the pathways GSK3β regulates that contribute to CIITA induction. One previous study in Raw264.7 cells found a requirement for GSK3β to activate STAT3 following IFNγ stimulation (51). While STAT1, and not STAT3, is thought to control the majority of CIITA induction, a minor role for STAT3 remained possible. To test the contribution of STAT1 and STAT3 to IFNγ-induced MHCII, we targeted both Stat1 and Stat3 with two independent sgRNAs to generate loss-of-function macrophages. As expected, when these cells were stimulated with IFNγ, Stat1 prevented the increase of MHCII surface expression. In contrast, neither sgRNA targeting Stat3 showed any difference in MHCII expression compared to non-targeting control (Figure S3C). Thus, while GSK3β may regulate STAT3 dependent pathways following IFNγ, a loss in STAT3 functionality does not explain the contribution of GSK3β to IFNγ-mediated MHCII induction. Together these results suggest that GSK3β, similarly to MED16, controls IFNγ-mediated MHCII expression upstream of the transcriptional induction of CIITA.

### GSK3α controls IFNγ-induced MHCII expression in the absence of GSK3β

Throughout the experiments above we observed that cells treated with CHIR99021 inhibited MHCII even more robustly than the Gsk3β KO cells. CHIR99021 not only inhibits GSK3β but also the paralog GSk3a (52). This led us to consider the role of GSK3a in IFNγ-mediated MHCII expression. While we did not observe enrichment for GSK3a in the screen (Figure 1D and Table S1), we could not exclude the possibility that GSK3a can play a secondary function during IFNγ-activation. Given the increased inhibition of MHCII with CHIR99021, we hypothesized that GSK3a can partially compensate for total loss of GSK3β, resulting in some remaining IFNγ-induced MHCII expression. To test this hypothesis, we treated NTC and Gsk3β KO macrophages with CHIR99021 or DMSO and quantified MHCII surface expression. Consistent with our previous results, we found robust inhibition of MHCII expression on NTC macrophages treated with CHIR99021 (Figure 4A and 4B). In support of a minor role for GSK3a, CHIR99021 treatment of Gsk3β KO macrophages further reduced surface MHCII expression after IFNγ-stimulation.

**Figure 4.**
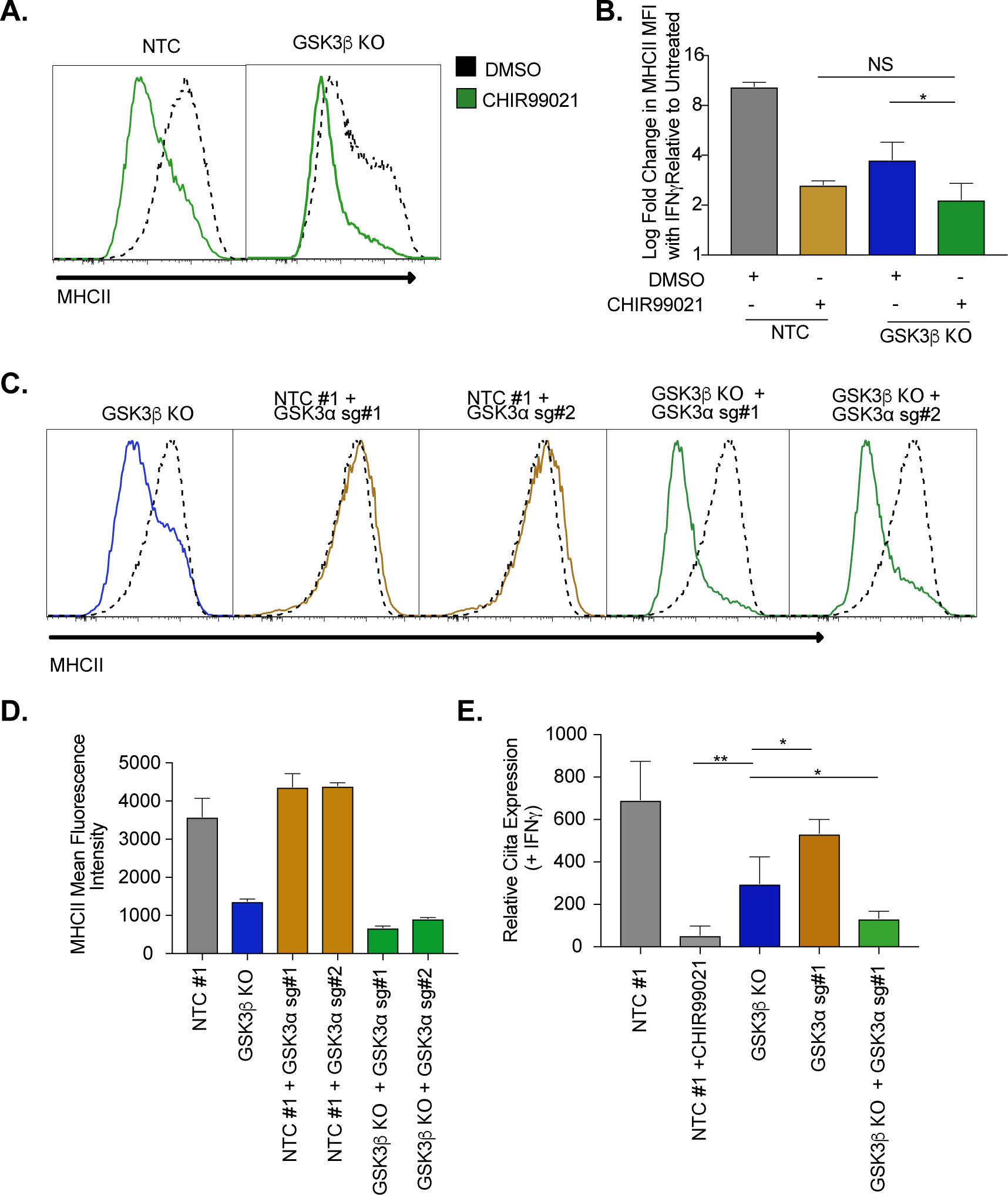
GSK3α controls IFNγ-mediate MHCII expression only in the absence of GSK3β. **(A)** NTC L3 cells or Gsk3β KO were treated with DMSO or 10uM CHIR99021 then left untreated or stimulated with IFNγ for 18 hours. MHCII surface expression was then quantified by Flow cytometry. A representative flow cytometry plot of DMSO treated IFNγ treated NTC cells (Dashed line) is overlaid with either NTC **(Left)** or Gsk3β KO **(Right)** treated with CHIR99021. The mean log fold change in MFI (IFNγ-treated/untreated) was quantified from three biological replicates. These results are representative of three independent experiments. **(C)** L3 cells or Gsk3β KO transduced with the indicated sgRNAs were treated with IFNγ and 18 hours later the surface levels of MHCII were quantified by flow cytometry. A representative histogram of NTC cells (dotted line) is overlaid with cells targeted with the indicated sgRNA (solid line) after 18 hours of IFNγ treatment. In **(D)** the mean fluorescence intensity of surface MHCII was quantified from 3 biological replicates from this experiment. **(E)** L3 cells or Gsk3β KO transduced with the indicated sgRNAs were treated with IFNγ and 18 hours later RNA was isolated and the expression of CIITA was quantified relative to GAPDH using qRT-PCR. Results are from three independent wells and are representative of two independent experiments. ***p<.001, **p<.01, *p<.05 by one-way ANOVA with a Tukey correction test.

To exclude the possibility of CHIR99021 off-target effects we next targeted Gsk3a genetically. To enable positive selection of a second sgRNA, we engineered vectors in the sgOpti background with distinct resistance markers for bacterial and mammalian selection that facilitated multiplexed sgRNA cloning (see materials and methods) (53). These vectors could be used to improve knockout efficiency when targeting a gene with multiple sgRNAs or target multiple genes simultaneously (Figure S4A). We targeted Gsk3a with two unique sgRNAs in either NTC or Gsk3β KO macrophages and stimulated the cells with IFNγ. Cells targeting Gsk3a alone upregulated MHCII expression similarly to NTC control cells (Figure 4C and 4D). In contrast, targeting Gsk3a in Gsk3β KO macrophages led to a further reduction of MHCII surface expression, similar to what was observed with CHIR99021 treatment. This same trend was observed when we examined CIITA mRNA expression after IFNγ-activation (Figure 4E). Therefore, blocking both GSK3a/b function results in a more severe reduction of IFNγ-stimulated MHCII expression in macrophages than GSK3β alone. Taken together, we conclude that GSK3a regulates IFNγ-mediated CIITA induction only in the absence of the GSK3β.

### GSK3β and MED16 function through distinct mechanisms to control IFNγ-mediated CIITA expression

Since the loss of either MED16 or GSK3β reduced IFNγ-mediated CIITA transcription, it remained possible that these two genes control MHCII expression through the same regulatory pathway. While Med16 KO macrophages are greatly reduced in IFNγ-mediated MHCII induction, there remains a small yet reproducible increase in MHCII surface expression. We determined if this effect on MHCII expression after IFNγ-activation required GSK3 activity by treating with CHIR99021. While DMSO-treated Med16 KO cells showed a reproducible 2-3 fold increase in MHCII expression after IFNγ stimulation, CHIR99021 treated Med16 KO cells showed no change whatsoever (Figure 5A and 5B). These results led us to hypothesize that MED16 and GSK3β control IFNγ-mediated CIITA induction and MHCII expression through independent mechanisms.

**Figure 5.**
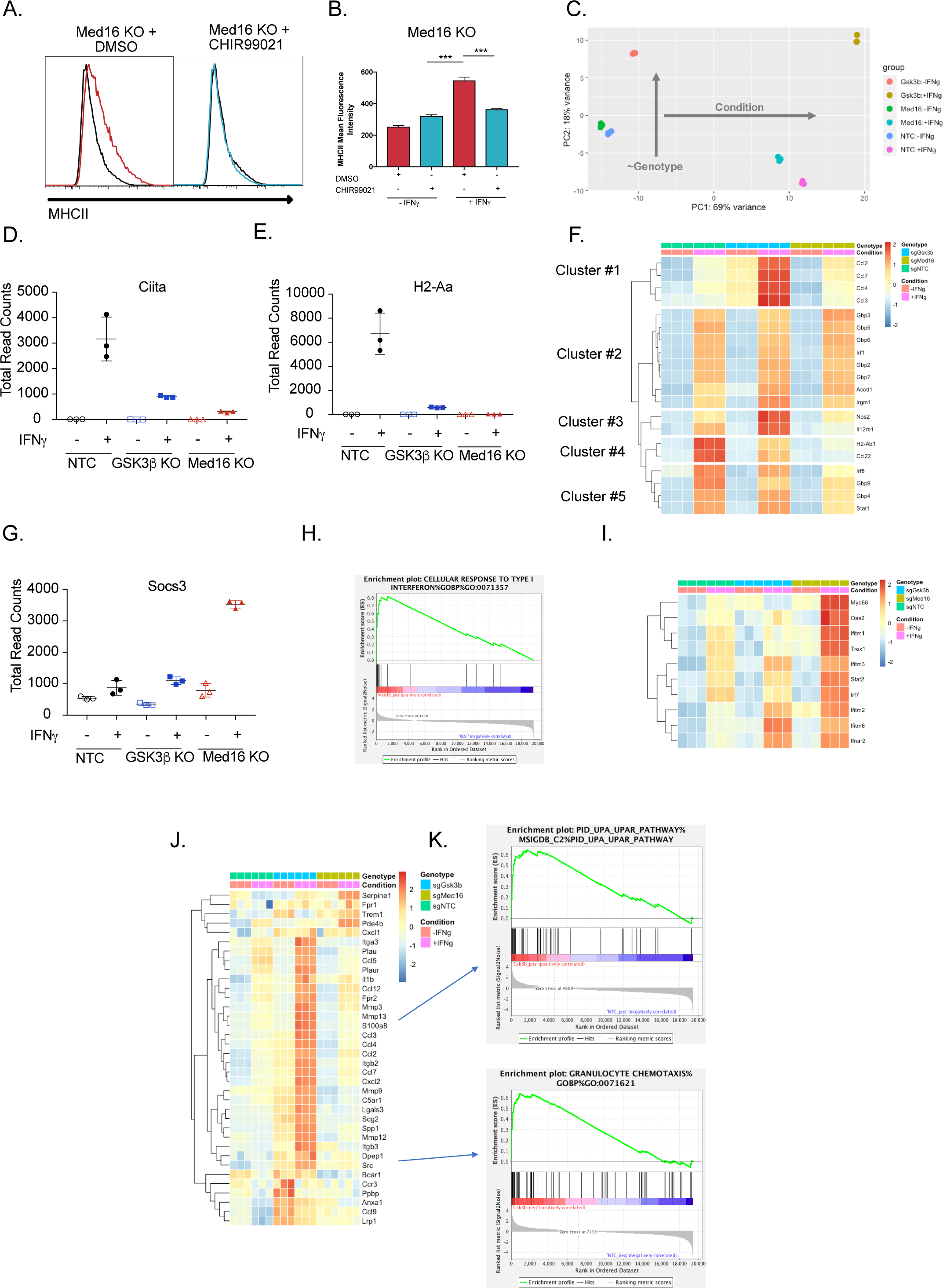
Transcriptomic analysis reveals distinct regulatory mechanisms of IFNγ signaling mediated by MED16 and GSK3β. **(A)** Med16 KO cells were treated with DMSO or CHIR99021 then left untreated or stimulated with IFNγ overnight. The following day MHC II cell surface expression was determined by flow cytometry. Shown is a representative histogram with the indicated treatment in untreated (Grey/Black line) or IFNγ-treated (Colored line) cells. **(B)** The quantification of the MFI of MHCII from four biological replicates. ***p<.001 by one-way ANOVA with Tukey correction. **(C)** The Global transcriptomes of NTC, Gsk3β KO and Med16KO was determined in the presence and absence of IFNγ-stimulation for 18 hours by RNA sequencing. Shown is the principal component analysis of the transcriptomes from three biological replicates for each condition. **(D)** Dotplot showing the normalized read counts for CIITA and **(E)** H2-Aa **(F)** Shown is a heatmap showing the relative expression (log normalized, row-scaled) of the most varied 20 genes involved in the cellular response to type II interferon (Gene Ontology GO:0071346). **(G)** Shown is a Dotplot visualizing the normalized counts of the type I IFN signature Socs3 from all RNAseq conditions. Clustering was used to **(H)** Significant gene sets from Med16 KO cells that were uniquely regulated from the RNAseq dataset were analyzed by gene set enrichment analysis (GSEA) then subjected to Leading Edge analysis, which identified a significant enrichment of the cellular responses to type I interferons (normalized enrichment score 2.81, FDR<0.01). **(I)** Shown is a heatmap demonstrating the relative expression of the type I interferon signature identified in IFNγ-stimualted Med16 KO macrophages from the RNAseq analysis. (J) Shown is a heatmap demonstrating the relative expression of unique differentially expressed genes from the Gsk3β KO in the presence (Top) and absence (Bottom) of IFNγ-stimulation. (K) These differentially expressed genes were used in GSEA to identify Leading Edge networks that are specific to Gsk3β KO cells. (Top) Shown is the leading-edge analysis of the UPAR pathway that was identified from IFNγ-stimulated Gsk3β KO cells. (Bottom) Shown is the leading-edge analysis of the Granulocyte chemotaxis pathway that was identified as differentially regulated in resting Gsk3β KO cells.

To test this hypothesis, we compared the transcriptional profiles of Med16 KO and Gsk3β KO cells to NTC cells by performing RNAseq on cells that were left untreated or were stimulated with IFNγ (See materials and methods). Principal component analysis of these 6 transcriptomes revealed distinct effects of IFNγ-stimulation (“condition”; PC1) and genotype (PC2) gene expression (Figure 5C). Both Med16 and Gsk3β knockout macrophages had distinct transcriptional signatures in the absence of cytokine stimulation, which were further differentiated with IFNγ-stimulation. The PCA analysis suggested that MED16 and GSK3β control distinct transcriptional networks in macrophages following IFNγ-activation.

Transcriptional analysis confirmed a critical role of GSK3β and MED16 in regulating IFNγ-dependent CIITA and MHCII expression in macrophages compared to NTC controls (Figure 5D and 5E). However, the extent to which MED16 or GSK3β controlled the overall response of macrophages to IFNγ remained unclear. To directly assess how MED16 and GSK3β regulate the general response to IFNγ, we queried IFNγ-regulated genes from our dataset that are annotated as part of the cellular response to IFNγ stimulation (GeneOntology:0071346). Hierarchical clustering found that, of the 20 most induced IFNγ-regulated transcripts, the expression of 8 were unaffected by loss of either GSK3β and MED16 (Figure 5F, Cluster 2). Importantly, these genes included a major regulator of the IFNγ response, IRF1, as well as canonical STAT1-target genes (GBP2, GBP3, GBP5, GBP6 and GBP7). This suggests that neither GSK3β nor MED16 are global regulators of the IFNγ response in macrophages, but rather are likely to exert their effect on particular genes at the level of transcription or further downstream. In contrast, only two genes, out the top 20 IFNγ-regulated genes, were similarly reduced in both Med16 KO and Gsk3β KO cells (Cluster 4), one of which was H2-Ab1. This shows that while GSK3β and MED16 both regulate IFNγ-mediated MHCII expression, they otherwise control distinct aspects of the IFNγ-mediated response in macrophages. The remaining clusters from this analysis showed specific changes in either Med16 KO or Gsk3β KO cells. Clusters 1 and 3 showed a subset of genes that were more robustly induced in Gsk3β KO cells compared to NTC and Med16 KO cells. These genes included NOS2, IL12RB1 and chemokines CCL2, CCL3, CCL4, and CCL7. In contrast, Cluster 5 showed a subset of genes that were reduced only in macrophages lacking MED16, including IRF8 and STAT1; as these effects were modest, and did not reach statistical significance, they may be suggestive of an incomplete positive feedforward in which MED16 plays a role. Further stringent differential gene expression analysis (FDR<0.05, absolute LFC>1) of the IFNγ-stimulated transcriptomes identified 69 and 90 significantly different genes for MED16 and GSK3β respectively. Of these differentially expressed genes (DEGs), eight non-MHCII genes were shared between MED16 and GSK3β, including five genes that are involved in controlling the extracellular matrix (MMP8, MMP12, TNN, and CLEC12a). Taken together these results suggest that while MED16 and GSK3β both regulate IFNγ-mediated CIITA and MHCII expression in macrophages, they otherwise control distinct regulatory networks in response to IFNγ.

We next used the transcriptional dataset to understand what aspects of IFNγ-mediated signaling MED16 and GSK3β specifically control. To resolve the transcriptional landscape of Med16 KO macrophages and to understand the specific effect that MED16 loss has on the host response to IFNγ, we analyzed the DEGs for upstream regulators whose effects would explain the observed gene expression signature. The analysis correctly predicted a relative inhibition on IFNγ signaling compared to NTC due to the muted induction of CIITA, H2-Ab1 and CD74. This analysis also identified signatures of IL-10, STAT3, and PPARg activation that included SOCS3 induction and PTGS2 downregulation (Figure 5G and Figure S5A and S5B). As the DEG analysis relied on a stringent threshold that filtered the great majority of the transcriptome from analysis, we sought to incorporate a more comprehensive analysis capable of capturing genes with more modest effects based on pathway enrichment. To this end, we performed gene set enrichment analysis (GSEA) using a ranked gene list derived from the differential gene expression analysis (54). Of the ∼10,000 gene sets tested, 11 sets were enriched for NTC + IFNγ and 76 for MED16 + IFNγ (FDR<0.1). To reduce pathway redundancy and infer biological relevance from the gene sets, we consolidated the signal into pathway networks (Figure S5C), and observed a significant enrichment for genes involved in xenobiotic and steroid metabolism, including many cytochrome p450 family members and glutathione transferases. We also observed an elevated type I interferon transcriptional response in Med16 KO cells stimulated with IFNγ that included components of IFNa/b signal transduction (IFNAR2), transcription factors (STAT2, IRF7) and antiviral mediators (OAS2, IFITM1, IFITM2, IFITM3, IFITM6) (Figure 5H and 5I). Thus, MED16 is a critical regulator of the overall interferon response in macrophages.

We next examined the regulatory networks that were specifically controlled by GSK3β. As observed by the initial PCA (Fig5C), the transcriptional landscape of GSK3β deficient macrophages was altered in unstimulated cells. We hypothesized that these widespread differences may alter cellular physiology and explain, in part, the varied responsiveness of Gsk3β KO cells to IFNγ. DEG analysis of unstimulated macrophages identified 284 differentially expressed genes due to GSK3β loss. Functional enrichment by STRING identified 3 major clusters that included dysregulation of chemokines, cell surface receptors, growth factor signaling, and cellular differentiation (FigS5D). We next examined the response of Gsk3β KO macrophages following IFNγ stimulation. GSEA identified a strong enrichment for chemotaxis and extracellular matrix remodeling pathways including several integrin subunits and matrix metalloproteinase members. These results suggest that GSK3β is an important regulator of both macrophage homeostasis and the response to IFNγ. Altogether the global transcriptional profiling suggests that while MED16 and GSK3β are both critical regulators of IFNγ-mediated MHCII expression, they each control distinct aspects of the macrophage response to IFNγ.

### Loss of MED16 or GSK3 inhibits macrophage-mediated CD4^+^ T cell activation

While the data to this point suggested that MED16 and GSK3β control the IFNγ-mediated induction of MHCII, in addition to distinct aspects of the IFNγ-response, it remained unclear how loss of GSK3β or MED16 in macrophages altered the activation of CD4^+^ T cells. To test this, we optimized an *ex vivo* T cell activation assay with macrophages and TCR-transgenic CD4^+^ T cells (NR1 cells) that are specific for the *Chlamydia trachomatis* antigen Cta1 (55). Resting NR1 cells were added to non-targeting control macrophages that were untreated, IFNγ stimulated, Cta1 peptide-pulsed, or IFNγ-stimulated and Cta1 peptide-pulsed. Five hours later, we harvested T cells and used intracellular cytokine staining to identify IFNγ producing cells by flow cytometry. Only macrophages that were treated with IFNγ and pulsed with Cta1 peptide were capable of stimulating NR1 cells to produce IFNγ (Figure 6A-6C). Additionally, when Rfx5 deficient macrophages were pulsed with peptide in the presence and absence of IFNγ, we observed limited IFNγ production by NR1 cells in both conditions suggesting this approach is peptide-specific and sensitive to macrophage MHCII surface expression.

**Figure 6.**
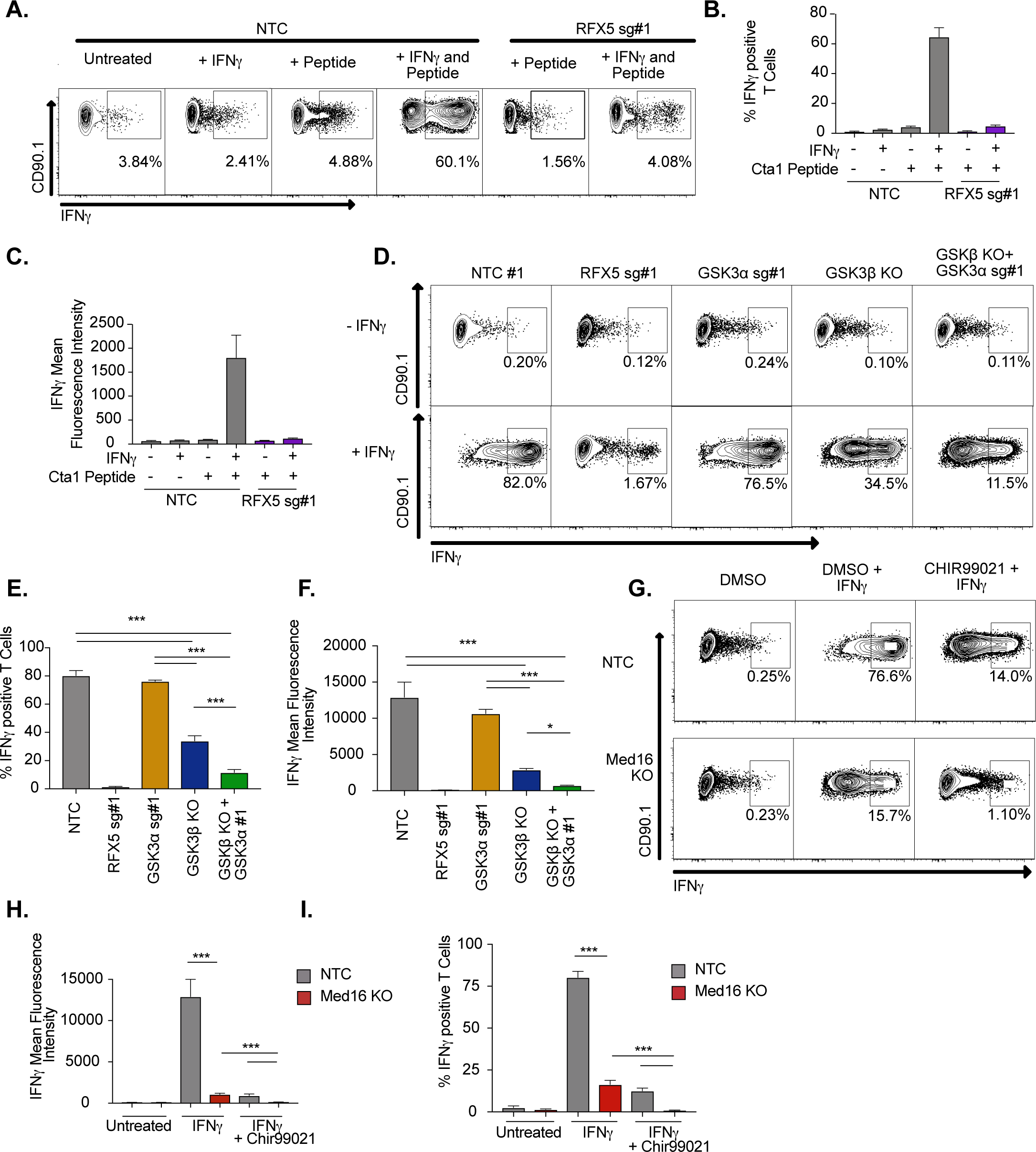
IFNγ-stimulated macrophages require MED16 or GSK3 to activate CD4^+^ T cells. **(A)** Macrophages were left untreated, treated with 10ng/ml IFNγ overnight, 5µM peptide for 1 hour or both IFNγ and peptide as indicated. TCR-transgenic NR1 CD4^+^ T cells specific for the peptide Cta1 from *Chlamydia trachomatis* were then added to L3 macrophages of the indicated genotypes at a 1:1 ratio. 4 hours after the addition of T cells, NR1 cells were harvested and the number of IFNγ-producing CD4^+^ T cells was quantified by intracellular staining and flow cytometry. Shown is a representative flow cytometry plot gated on live/CD4^+^ cells. Gates for IFNγ^+^ T cells were determined using an isotype control antibody. **(B)** The percent of live CD4^+^ T cells producing IFNγ and **(C)** the MFI of IFNγ production by live CD4^+^ T cells was quantified from triplicate samples. These results are representative of three independent experiments. **(D)** L3 cells targeted with the indicated sgRNAs were left untreated or treated overnight with IFNγ then pulsed with Cta1 peptide for 1 hour. NR1 cells were then added at a 1:1 ratio and 4 hours later NR1 cells were harvested and the number of IFNγ-producing CD4^+^ T cells was quantified by intracellular staining and flow cytometry. Shown is a representative flow cytometry plot gated on live/CD4^+^ cells. Gates for IFNγ^+^ T cells were determined using an isotype control antibody. **(E)** The percent of live CD4^+^ T cells producing IFNγ and **(F)** the MFI of IFNγ production by live CD4^+^ T cells was quantified from triplicate samples. These results are representative of three independent experiments. **(G)** NTC L3 cells or Med16 KO cells were left untreated or treated overnight with DMSO, IFNγ and DMSO or IFNγ and CHIR999021 then pulsed with Cta1 peptide for 1 hour. NR1 cells were then added at a 1:1 ratio and 4 hours after the addition of T cells, NR1 cells were harvested and the number of IFNγ-producing CD4^+^ T cells was quantified by intracellular staining and flow cytometry. Shown is a representative flow cytometry plot gated on live/CD4^+^ cells. Gates for IFNγ^+^ T cells were determined using an isotype control antibody. **(H)** The percent of live CD4^+^ T cells producing IFNγ and **(I)** the MFI of IFNγ production by live CD4^+^ T cells was quantified from triplicate samples. These results are representative of three independent experiments.

We next determined the effectiveness of macrophages lacking GSK3 components to activate CD4^+^ T cells. Macrophages deficient in GSK3α, GSK3β or GSK3α/β along with NTC and RFX5 controls were left untreated or stimulated with IFNγ for 16 hours, then all cells were pulsed with Cta1 peptide. Resting NR1 cells were then added and the production of IFNγ by NR1 cells from each condition was quantified by flow cytometry five hours later. In agreement with our findings on MHCII expression, loss of GSK3a did not inhibit the production of IFNγ by NR1 cells (Figure 6D-6F). In contrast, Gsk3β KO cells reduced the number of IFNγ^+^ NR1 cells over two-fold and reduced the mean fluorescence intensity of IFNγ production over 4-fold. Furthermore, macrophages deficient in GSK3a and GSK3β were almost entirely blocked in their ability to activate IFNγ production by NR1 cells. Thus, macrophages deficient in GSK3 function are unable to serve as effective antigen presenting cells to CD4^+^ T cells.

The *ex vivo* T cell assay was next used to test the effectiveness of Med16 KO macrophages as APCs. NR1 cells stimulated on IFNγ activated Med16 KO macrophages were reduced in the number of IFNγ^+^ T cells by 10-fold and the fluorescence intensity of IFNγ by 100-fold compared to NTC (Figure 6G-GI). Similar to what we observed with MHCII expression, there was a small yet reproducible induction of IFNγ^+^ NR1 cells incubated with IFNγ-activated Med16 KO macrophages. We hypothesized that inhibition of GSK3 and MED16 simultaneously would eliminate all NR1 activation on macrophages. Treatment of Med16 KO macrophages with CHIR99021 prior to IFNγ-stimulation and T cell co-incubation, eliminated the remaining IFNγ production by NR1 cells seen in the DMSO treated Med16 KO condition. Altogether these results show that GSK3β and MED16 are critical regulators of IFNγ mediated antigen presentation in macrophages and their loss prevents the effective activation of CD4^+^ T cells.

## Discussion

IFNγ-mediated MHCII is required for the effective host response against infections. Here, we used a genome-wide CRISPR library in macrophages to globally examine mechanisms of IFNγ-inducible MHCII expression. The screen correctly identified major regulators of IFNγ-signaling, highlighting the specificity and robustness of the approach. In addition to known regulators, our analysis identified many new positive regulators of MHCII surface expression. While we validated only a subset of these candidates, the high rate of validation suggests many new regulatory mechanisms of IFNγ-inducible MHCII expression in macrophages. While the major pathways identified from the candidates in CRISPR screen were related to IFNγ-signaling, we also identified an important role for other pathways including the mTOR signaling cascade. Within the top 100 candidates of the screen several genes related to metabolism and lysosome function including LAMTOR2 and LAMTOR4 were found. Given the known effects of IFNγ in modulating host metabolism, these results suggest that the metabolic changes following IFNγ-activation of macrophages is critical for key macrophage functions including the surface expression of MHCII (56). In addition, we found the small lysosome associated GTPase Arl8a is an important regulator of IFNγ-mediated MHCII surface expression. Interestingly, a paralog, Arl8b, was previously described as a regulator of MHCII and CD1d, by controlling lysosomal function (57, 58). Future studies will need to dissect the metabolism specific mechanisms that macrophages use to control the IFNγ response, including the regulation of MHCII.

In this study, we focused our follow up efforts from validated candidates on genes that might control MHCII transcriptional regulation. We identified MED16 and GSK3β as strong regulators of IFNγ-mediated CIITA induction. Using global transcriptomics we found that loss of either MED16 or GSK3β in macrophages inhibited subsets of IFNγ-mediated genes including MHCII. Importantly, the evidence here strongly supports a model where MED16 and GSK3β control IFNγ-mediated MHCII expression through distinct mechanisms (Figure 7). Our results uncover previously unknown regulatory control of CIITA-mediated expression that is biologically important to activate CD4^+^ T cells.

**Figure 7.**
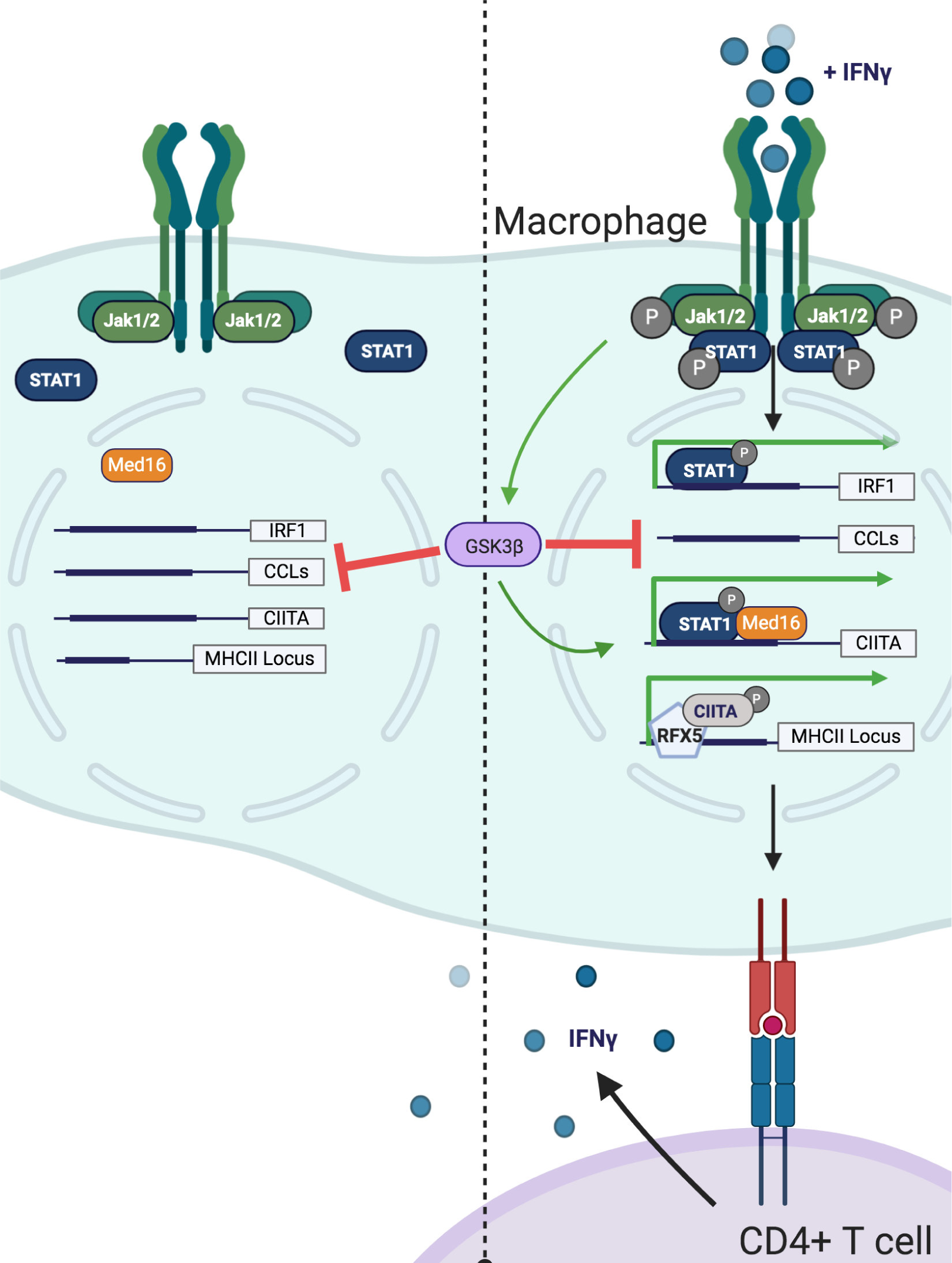
Model of GSK3β- and Med16-mediated control of IFNγ-activated MHCII expression. Shown is a model of how GSK3β and MED16 regulate IFNγ-mediated MHCII expression. In the absence of IFNγ **(Left)** GSK3β controls the transcription of many macrophage genes related to inflammation such as CCLs. In contrast, Med16 KO cells shows minimal transcriptional changes in resting macrophages. Additionally, IFNγ-mediated gene expression is low. Following the activation of macrophages with IFNγ **(Right)**, STAT1 becomes phosphorylated and translocates to the nucleus to drive gene transcription. The IFNγ-mediated induction of IRF1 does not require either GSK3β or MED16. While GSK3β continues to negatively regulate inflammatory genes like CCLs it also positively regulates the transcriptional activation of CIITA following IFNγ-activation. Through a parallel but distinct mechanism, IFNγ-mediated induction of CIITA also requires MED16 function. The expression of CIITA then recruits other transcription factors such as RFX5 to the MHCII locus where it induces the expression of MHCII, which allows for the activation of CD4^+^ T cells.

MED16 is a subunit of the mediator complex that is critical to recruit RNA polymerase II to the transcriptional start site (42). While the mediator complex can contain over 20 unique subunits and globally regulate gene expression, individual mediator subunits control distinct transcriptional networks by interacting with specific transcription factors (42, 44). Our data shows that MED16 is uniquely required among the mediator complex for IFNγ-mediated MHCII expression. How MED16 controls CIITA expression remains an open question. One recent study showed that MED16 controls NRF2 related signaling networks that respond to oxidative stress (59). A major finding of our MED16 transcriptional analysis was the identification of several metabolic pathways involved in oxidative stress and xenobiotics. Given the previous work that described how oxidative stress and the NRF2 regulator Keap1 regulated IFNγ-mediated MHCII expression in human melanoma cells, NRF2 regulation and redox dysregulation could explain a possible mechanism for MED16 control of MHCII (17). Intriguingly, the effect of MED16 loss was negligible on many STAT1 and IRF1 targets, and, in fact, resulted in a type I interferon gene signature. Whether this signature is causative of or secondary to the dysregulated response to type II interferons remains unknown.

Previous studies showed that CDK8, a kinase that can associate with the mediator complex, controls a subset of IFNγ-dependent gene transcription (60). However, our results strongly support a model where MED16 acts independently of CDK8. Not only was CDK8 not identified in the initial CRISPR screen, but our transcriptional profiling showed that the major IFNγ-dependent genes controlled by CDK8, TAP1 and IRF1, remain unchanged in Med16 KO macrophages. Thus, understanding what transcription factors MED16 interacts with in the future will be needed to fully determine the mechanisms of MED16-dependent transcription and its control over CIITA and IFNγ-mediated gene expression.

While we hypothesize that MED16 directly controls CIITA transcription, GSK3 likely regulates MHCII through signaling networks upstream of transcription initiation. GSK3a and GSK3β are multifunctional kinases that regulate diverse cellular functions including inflammatory and developmental cascades (43). Our studies found that loss of GSK3β but not GSK3a blocked efficient IFNγ-mediated MHCII expression. However, ours results suggest that even though GSK3a is not a primary regulator of IFNγ-mediated MHCII expression, it can partially compensate for the loss of GSK3β. Thus, GSK3a and GSK3β are partially redundant in their control of IFNγ-mediated MHCII expression highlighting the interlinked regulation of MHCII. This finding supports using genetic interactions studies in the future to fully understand the IFNγ-mediated regulatory networks in macrophages.

Because GSK3 regulates a range of pathways, careful work will be needed to determine which GSK3 regulated networks are responsible for controlling CIITA expression. One major function of GSK3 is to modulate the activation of the Wnt signaling cascade (43). Inhibition or loss of GSK3 results in the constitutive stabilization of Beta-Catenin and TCF expression. If the constitutive activation of Beta-catenin and Wnt signaling prevents effective CIITA expression remains to be determined. Interestingly, another Wnt signaling pathway member FZD4 was identified in our screen as required for MHCII expression in our screen, supporting a possible role for Wnt in IFNγ-induced MHCII regulation. It is tempting to speculate that Wnt signaling balances IFNγ-induced activation, resulting in distinct MHCII upregulation between cells with different Wnt activation states. While there is data supporting interactions between Wnt pathways and Type I IFN during viral infections, this has not been explored yet in the context of IFNγ (61, 62).

Previous studies suggested that GSK3 controls IFNγ mediated STAT3 activation, LPS-mediated nitric oxide production, and IRF1 transcriptional activity but our results in macrophages clearly show these do not explain the requirement for GSK3-dependent MHCII expression (51, 63, 64). In contrast, we found no role for STAT3 in IFNγ-mediate MHCII expression and significantly higher expression of inducible Nitric Oxide Synthase in Gsk3β KO macrophages. In addition, we observed a significant increase in a number of chemokines that are critical to mobilizing cells to the site of infections. These results show that GSK3 is a central regulator of the balanced host response during infection, and that targeting GSK3 function is likely to make the host susceptible to disease. In line with this prediction, GSK3 was recently found to be co-opted by the *Salmonella enterica* serovar Typhimurium effector SteE to skew infected macrophage polarization and allow infection to persist (65, 66). Our results suggest another possible effect of targeting GSK3 may be the inefficient upregulation of MHCII on Salmonella-infected macrophages in response to IFNγ. While it is known that *Salmonella* and other pathogens including *M. tuberculosis* and *C. trachomatis*, modulate the expression of MHCII, the precise mechanisms underlying many of these virulence tactics remains unclear (27, 28). Our screening results provide a framework to test the contribution of each candidate MHCII regulator during infection with pathogens that target MHCII. These directed experiments would allow the rapid identification of possible host-pathogen interactions. It will be important to determine if augmenting specific MHCII pathways identified by our screen overcomes pathogen-mediated inhibition and induces robust MHCII expression to better activate CD4^+^ T cells and protect against disease.

Beyond infections, our dataset provides an opportunity to examine the importance of newly identified MHCII regulators in other diseases such as tumor progression and autoimmunity. Of course, MHCII is not the only surface marker that is targeted by pathogens and malignancy. Other important molecules including MHCI, CD40 and PD-L1 are induced by IFNγ stimulation and are targeted in different disease states (67–70). Employing our screening pipeline for a range of surface markers will identify regulatory pathways that are shared and unique at high resolution and provide insights into targeting these pathways therapeutically. Taken together, the tools and methods developed here identified new regulators of IFNy­ MHCII that will illuminate the underlying biology of the host immune response.

## METHODS

### Mice

C57BL/6J (stock no. 000664) were purchased from The Jackson Laboratory. NR1 mice were a gift of Dr. Michael Starnbach (55). Mice were housed under specific pathogen-free conditions and in accordance with the Michigan State University Institutional Animal Care and Use Committee guidelines. All animals used for experiments were 6–12 weeks of age.

### Cell culture

Macrophage cell lines were maintained in Dulbecco’s Modified Eagle Medium (DMEM; Hyclone) supplemented with 5% fetal bovine serum (Seradigm). Cells were kept in 5% CO2 at 37C. For HoxB8-conditionally immortalized macrophages, bone marrow from C57BL6/J mice was transduced with retrovirus containing estradiol-inducible HoxB8 then maintained in media containing 10% GM-CSF conditioned supernatants, 10% FBS and 10uM Beta-Estradiol as previously described (50). To generate BMDMs cells were washed 3x in PBS to remove estradiol then plated in 20% L929 condition supernatants and 10% FBS. 8-10 days later cells were plated for experiments as described in the figure legends.

### CRISPR Screen and Analysis

The mouse BRIE knockout CRISPR pooled library was a gift of David Root and John Doench (Addgene #73633) (36). Using the BRIE library, 4 sgRNAs targeting every coding gene in mice in addition to 1000 non-targeting controls (78,637 sgRNAs total) were packaged into lentivirus using HEK293T cells and transduced in L3 cells at a low multiplicity of infection (MOI <0.3) and selected with puromycin two days after transduction. Sequencing of the input library showed high coverage and distribution of the library (FigS1). We next treated the library with IFNγ (10ng/ml) and 24 hours later the cells were fixed and fluorescence activated cell sorting (FACS) was used to isolate the MHCII^high^ and MHCII^low^ bins. Bin size was guided by the observed phenotypes of positive control sgRNAs, such as RFX5, which were tested individually and to ensure sufficient coverage (>25x unselected library) in the sorted populations. Genomic DNA was isolated from sorted populations from two biological replicate experiments using Qiagen DNeasy kits. Amplification of sgRNAs by PCR was performed as previously described using Illumina compatible primers from IDT (36), and amplicons were sequenced on an Illumina NextSeq500.

Sequence reads were first trimmed to remove any adapter sequence and to adjust for p5 primer stagger. We used bowtie 2 via MAGeCK to map reads to the sgRNA library index without allowing for any mismatch. Subsequent sgRNA counts were median normalized to control sgRNAs in MAGeCK to account for variable sequencing depth. Control sgRNAs were defined as non-targeting controls as well as genes not-transcribed in our macrophage cell line as determined empirically by RNA-seq (Table S2). To test for sgRNA and gene enrichment, we used the ‘test’ command in MAGeCK to compare the distribution of sgRNAs in the MHCII^high^ and MHCII^low^ bins. Notably, we included the input libraries in the count analysis in order to use the distribution of sgRNAs in the unselected library for the variance estimation in MAGeCK.

### sgRNA cloning

sgOpti was a gift from Eric Lander & David Sabatini (Addgene plasmid #85681) (53). Individual sgRNAs were cloned as previously described (71). Briefly, annealed oligos containing the sgRNA targeting sequence were phosphorylated and cloned into a dephosphorylated and BsmBI (New England Biolabs) digested SgOpti (Addgene#85681) which contains a modified sgRNA scaffold for improved sgRNA-Cas9 complexing. A detailed cloning protocol is available in supplementary methods. To facilitate rapid and efficient generation of sgRNA plasmids with different selectable markers, we further modified the SgOpti vector such that the mammalian selectable marker was linked with a distinct bacterial selection. Subsequent generation of SgOpti-Blasticidin-Zeocin (BZ), SgOpti-Hygromycin-Kanamycin (HK), and SgOpti-G418-Hygromycin (GH) allowed for pooled cloning in which a given sgRNA was ligated into a mixture of BsmBI-digested plasmids. Successful transformants for each of the plasmids were selected by plating on ampicillin (SgOpti), zeocin (BZ), kanamycin (HK), or hygromycin (GH) in parallel. In effect, this reduced the cloning burden 4x and provided flexibility with selectable markers to generate near-complete editing in polyclonal cells and/or make double knockouts.

### Flow cytometry

Cells were harvested at the indicated times post-IFNγ stimulation by scrapping to ensure intact surface proteins. Cells were pelleted and washed with PBS before staining for MHCII. MHCII expression was analyzed on the BD LSRII cytometer or a BioRad S3E cell sorter. All flow cytometry analysis was done in FlowJo V9 or V10 (TreeStar)

### Chemical inhibitors

CHIR99021 (Sigma) was resuspended in DMSO at 10 mM stock concentration. DMSO was added at the same concentration to the inhibitors as a control. Cells were maintained in 5% CO_2_. Cells were stimulated with 6.25ng/ml of IFNγ (Biolegend) for the indicated times in each figure legend before analysis.

### Isolation of Knockout cells

Cells transduced with either MED16 or GSK3β sgRNAs were stimulated with IFNγ then stained for MHCII 24 hours later. Cells expressing low MHCII were then sorted using a BioRad S3e cell sorter and plated for expansion. Gene knockouts were confirmed by amplifying the genomic regions encoding either MED16 or GSK3β from each cell population in addition to NTC cells using PCR. PCR products were purified by PCR-cleanup Kit (Qiagen) and sent for Sanger Sequencing (Genewiz). The resultant ABI files were used for TIDE analysis to assess the frequency and size of indels in each population compared to control cells.

### RNA isolation

Macrophages were homogenized in 500uL of TRIzol reagent (Life Technologies) and incubated for 5 minutes at room temperature. 100uL of chloroform was added to the homogenate, vortexed, and centrifuged at 12,000 x g for 20 minutes at 4C to separate nucleic acids. The clear, RNA containing layer was removed and combined with 500uL of ethanol. This mixture was placed into a collection tube and protocols provided by the Zymo Research Direct-zol RNA extraction kit were followed. Quantity and purity of the RNA was checked using a NanoDrop and diluted to 5ng/uL in nuclease-free water.

### RNA-sequencing Analysis

To generate RNA for sequencing, macrophages were seeded in 6-well dishes at a density of 1 million cells/well. Cells were stimulated for 18 hours with IFNγ (Peprotech) at a final concentration of 6.25 ng/mL, after which RNA was isolated as described above. RNA quality was assessed by qRT-PCR as described above and by TapeStation (Aligent); the median RIN value was 9.5 with a ranger of 8.6 to 9.9. A standard library preparation protocol was followed to prepare sequencing libraries on poly-A tailed mRNA using the NEBNext® Ultra™ RNA Library Prep Kit for Illumina®. In total, 18 libraries were prepared for dual index paired-end sequencing on a HiSeq 2500 using a high-output kit (Illumina) at an average sequencing depth of 38.6e6 reads per library with > 93% of bases exceeding a quality score of 30. FastQC (v0.11.5) was used to assess the quality of raw data. Cutadapt (v2.9) was used to remove TruSeq adapter sequences with the parameters --cores=15 -m 1 -a AGATCGGAAGAGCACACGTCTGAACTCCAGTCA -A AGATCGGAAGAGCGTCGTGTAGGGAAAGAGTGT. A transcriptome was prepared with the rsem (v1.3.0) command rsem-prepare-reference using bowtie2 (v2.3.5.1) and the gtf and primary Mus musculus genome assembly from ENSEMBL release 99. Trimmed sequencing reads were aligned and counts quantified using rsem-calculate-expression with standard bowtie2 parameters; fragment size and alignment quality for each sequencing library was assessed by estimating the read start position distribution (RSPD) via --estimate-rspd. Gene counts as determined by rsem were used as input for differential expression analysis in DESeq2 according to standard protocols. Briefly, counts were imported using tximport (v1.16.0) and differential expression was performed with non-targeting control (“NTC”) and unstimulated (“Condition A”) as reference levels for contrasts. For visualization via PCA, a variance stabilizing transformation was performed in DESeq2. Pathway enrichment utilized R packages gage and fgsea or Ingenuity Pathway Analysis (Qiagen). Gene-set enrichment analysis (GSEA) was performed utilized gene rank lists as calculated from defined comparisons in DeSeq2 and was inclusive of gene sets comprised of 10-500 genes that were compiled and made available by the Bader lab (72). Pathway visualization and network construction was performed in CytoScape 3.8 using the apps STRING and EnrichmentMap. Pathway significance thresholds were set at an FDR of 0.1 unless specified otherwise.

### Quantitative real time PCR

PCR amplification of the RNA was completed using the One-step Syber Green RT-PCR Kit (Qiagen). 25ng of total RNA was added to a master mix reaction of the provided RT Mix, Syber green, gene specific primers (5uM of forward and reverse primer), and nuclease-free water. For each biological replicate (triplicate), reactions were conducted in technical duplicates in 96-well plates. PCR product was monitored using the QuantStudio3 (ThermoFisher). The number of cycles needed to reach the threshold of detection (Ct) was determined for all reactions. Relative gene expression was determined using the 2^-ddCT method. The mean CT of each experimental sample in triplicate was determined. The average mean of glyceraldehyde 3-phosphate dehydrogenase (GAPDH) was subtracted from the experimental sample mean CT for each gene of interest (dCT). The average dCT of the untreated control group was used as a calibrator and subtracted from the dCT of each experimental sample (ddCT). 2^-ddCT shows the fold change in gene expression of the gene of interest normalized to GAPDH and relative to to untreated control (calibrator).

### T cell activation assays

CD4^+^ T cells were harvested from the lymph nodes and spleens of naive NR1 mice and enriched with a mouse naïve CD4 negative isolation kit (BioLegend) following the manufacturer’s protocol. CD4^+^ T cells were cultured in media consisting of RPMI 1640 (Invitrogen), 10% FCS, l-glutamine, HEPES, 50 μM 2-ME, 50 U/ml penicillin, and 50 mg/ml streptomycin. NR1 cells were activated by coculture with mitomycin-treated splenocytes pulsed with 5 μM Cta1_133–152_ peptide at a stimulator/T cell ratio of 4:1. Th1 polarization was achieved by supplying cultures with 10 ng/ml IL-12 (PeproTech, Rocky Hill, NJ) and 10 μg/ml anti–IL-4 (Biolegend) One week after initial activation resting NR1 cells were co-incubated with untreated or IFNγ-treated macrophages of different genotypes, that were or were not pulsed with Cta1 peptide. Six hours following co-incubation NR1 cells were harvested and stained for intracellular IFNγ (BioLegend) using an intracellular cytokine staining kit (BioLegend) as done previously. Analyzed T cells were identified as live, CD90.1^+^ CD4^+^ cells.

### Statistical Analysis and Figures

Statistical analysis was done using Prism Version 7 (GraphPad) as indicated in the figure legends. Data are presented, unless otherwise indicated, as the mean ^+^/− the standard deviation. Figures were created in Prism V7 or were created with BioRender.com

## Acknowledgements

We would like thank members of the Sassetti, Abramovitch and Olive labs for critical feedback and input throughout the project. We thank Dr. Robert Abramovitch for critical reading of the manuscript. We thank the flow cytometry core at MSU and UMMS for their help in all experiments requiring flow cytometry. We would also like to thank Dr. Michael Starnbach for the gift of the NR1 mice. This work was supported by startup funding to AJO provided by Michigan State University, support from the Arnold O. Beckman Postdoctoral fellowship to AJO and grants from the NIH (AI146504, AI132130).

---

**Figure S1.**
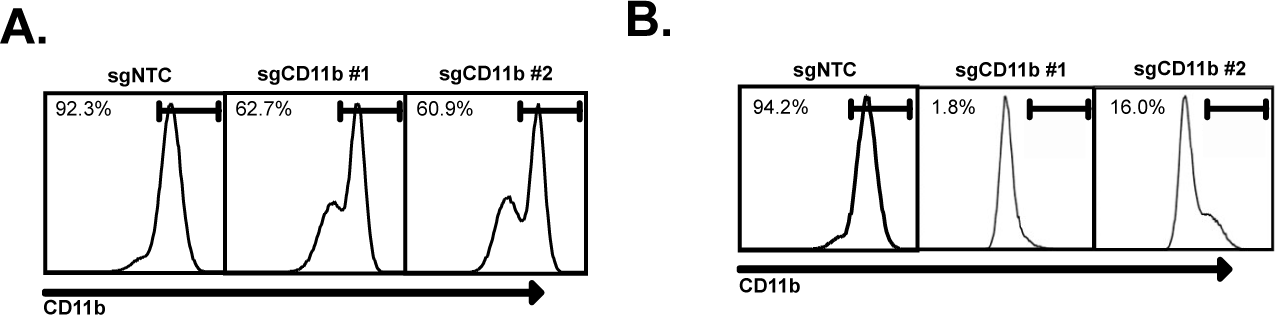
Optimization of CRISPR-Cas9 editing in iBMDMs. Immortalized C57BL6/J macrophages were transduced with lentivirus expressing Cas9 then selected using Hygromycin. Polyclonal transductants were then transduced with a second lentivirus encoding two different sgRNAs targeting CD11b or a non-targeting control then selected with puromycin. **(A)** Transductants were then stained for surface CD11b one-week later and analyzed by flow cytometry. Shown is a representative histogram of CD11b from the polyclonal Cas9 line. **(B)** Single cell clones were isolated from the polyclonal Cas9 line by limiting dilution. One clone, clone L3 was transduced with two different sgRNAs targeting CD11b or a non-targeting control then selected with puromycin. Transductants were then stained for surface CD11b one-week later and analyzed by flow cytometry. Shown is a representative histogram of CD11b from the L3 Cas9 clone.

**Figure S2.**
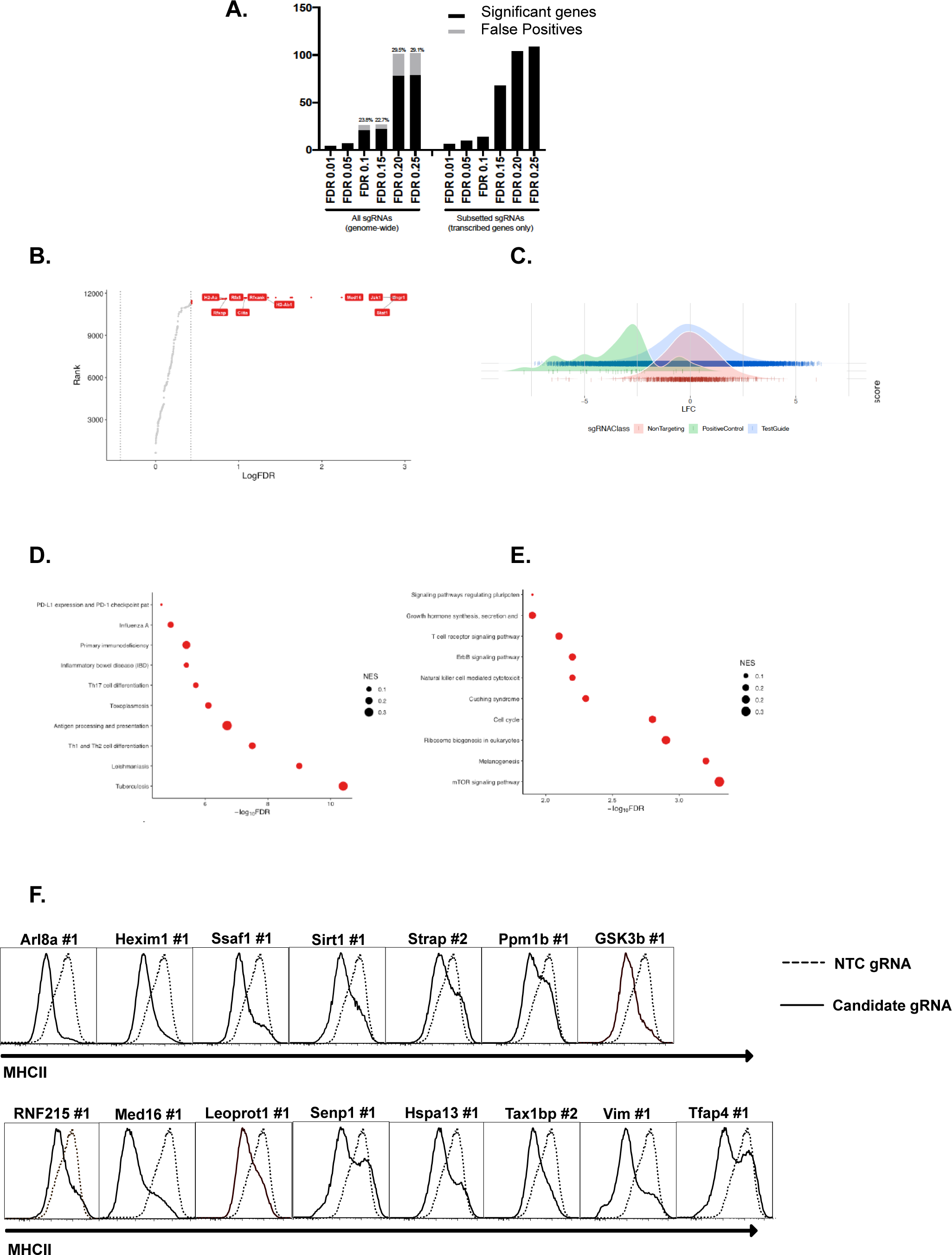
Adaptations to the MAGeCK analysis pipeline identifies high confidence regulators of IFNγ-mediated MHCII expression following a Genome-wide CRISPR Cas9 screen. **(A)** At the selected alpha cutoff of 0.025, the number of significant genes by FDR level and the number of false positives (gray bar within black bar) when using all guides; proportion of significant genes that were false positives annotated above each bar (left half). This analysis was repeated using only transcribed genes as determined by RNAseq analysis (Table S2). **(B)** Genes that passed quality filtering and were expressed within L3 cells at the RNA level as determined by transcriptomics were ranked by the FDR as determined in MAGeCK. Highlighted in Red are the top hits many of which are within the canonical IFNγ-signaling and MHCII expression pathway. Vertical dotted lines indicate genes that are below a calculated FDR of .2 and include the follow-up candidates Med16 and GSK3β. **(C)** The distribution of sgRNAs based on the Log2-Fold Change as determined by MAGeCK from three groups of sgRNA targets; Positive controls (Known IFNγ/MHCII pathway), Test Guides (all targeting guides in BRIE library) and non-targeting controls (∼1000 included in BRIE library) are shown. Positive control sgRNAs are left shifted compared to negative controls indicating an enrichment in the MHCII^low^ population of these sgRNAs. **(D)** Significant genes from the genome-wide screen were used to identify enriched pathways from the KEGG pathway database. Shown are the top 10 enriched pathways from the screen results indicating a significant enrichment of IFNγ-related pathways ranked by FDR and the size of the circle indicates the Normalized Enrichment Score. **(E)** To identify new pathways unrelated to IFNγ signaling KEGG pathway enrichment was repeated with the top 11 genes related to IFNγ removed from the query list. Shown are the top 10 pathways that were identified by KEGG ranked by FDR and the size of the circle indicates the Normalized Enrichment Score. **(F)** The L3 clone was transduced with the indicated sgRNAs for candidates in the top 100 candidates from the CRISPR-Cas9 screen. All cells were treated with 10ng/ul of IFNγ for 24 hours then were analyzed by flow cytometry. Shown are representative flow cytometry plots from the data quantified in Figure 1E. The results are representative of at least two independent experiments. *p<.05 **p<.01 ***p<.001 by one-way ANOVA compared to the mean of NTC1 and NTC2 using Dunnets multiple comparison test.

**Figure S3.**
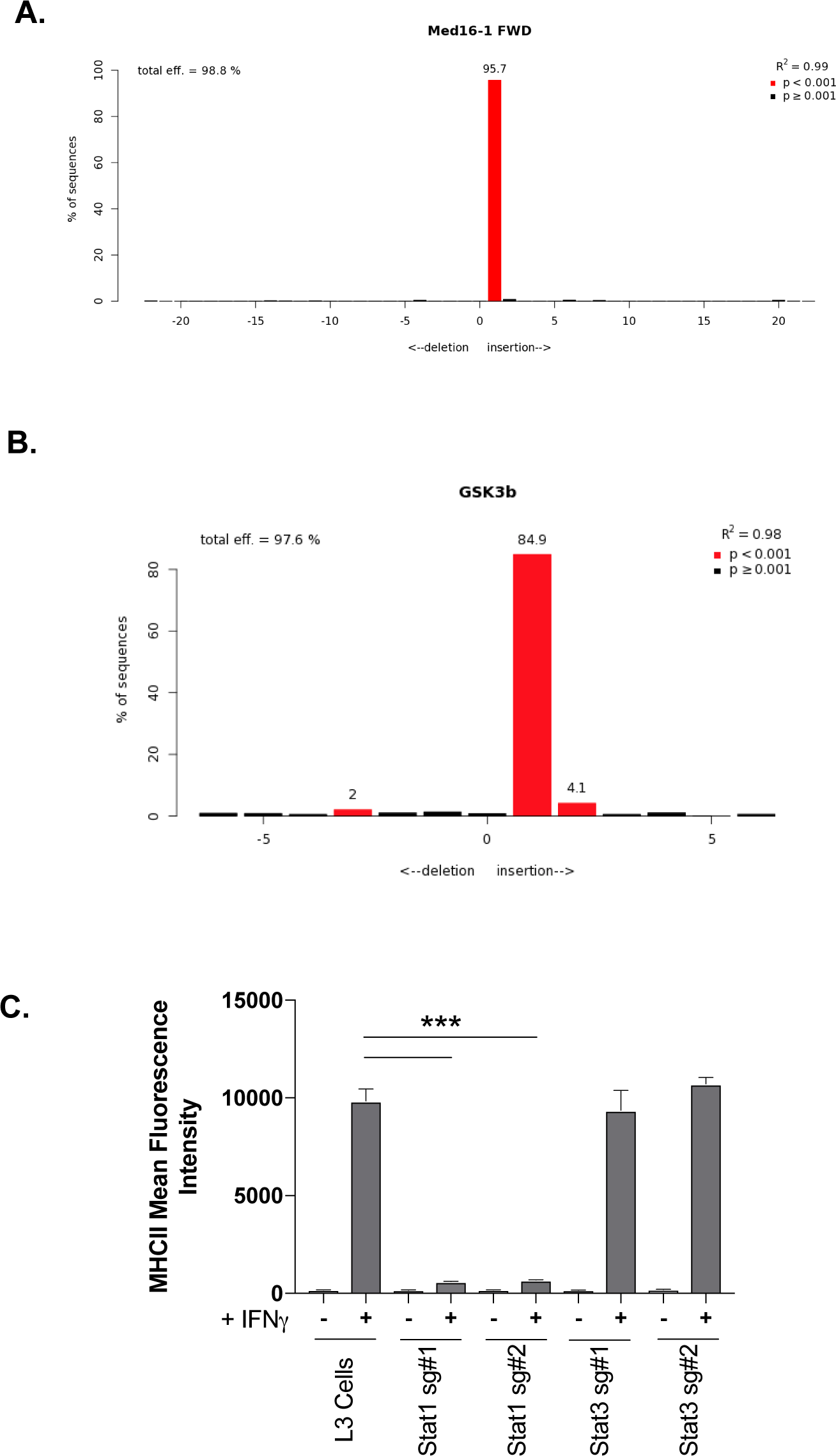
Confirmation of KO lines using TIDE analysis. Genomic DNA was isolated from NTC, Med16 KO and Gsk3β KO cells and the PCR was used to amplify the region encoding either Med16 or GSK3β. TIDE analysis was used to quantify the editing efficiency of the indels in each cell line using trace plots following Sanger sequencing. Shown is the TIDE analysis profile indicating the percent editing efficiency for (A) Med16 KO and (B) Gsk3β KO cells compared to NTC control cells. (C) L3 cells and cells transduced with sgRNAs targeting either Stat1 or Stat3 were left untreated or were stimulated with IFNγ for 18 hours. The surface levels of MHCII were then quantified by flow cytometry and the mean fluorescence intensity was determined from triplicate samples. These results are representative of two independent experiments. ***p<.001 by one-way ANOVA compared using Dunnets multiple comparison test compared to L3 controls.

**Figure S4.**
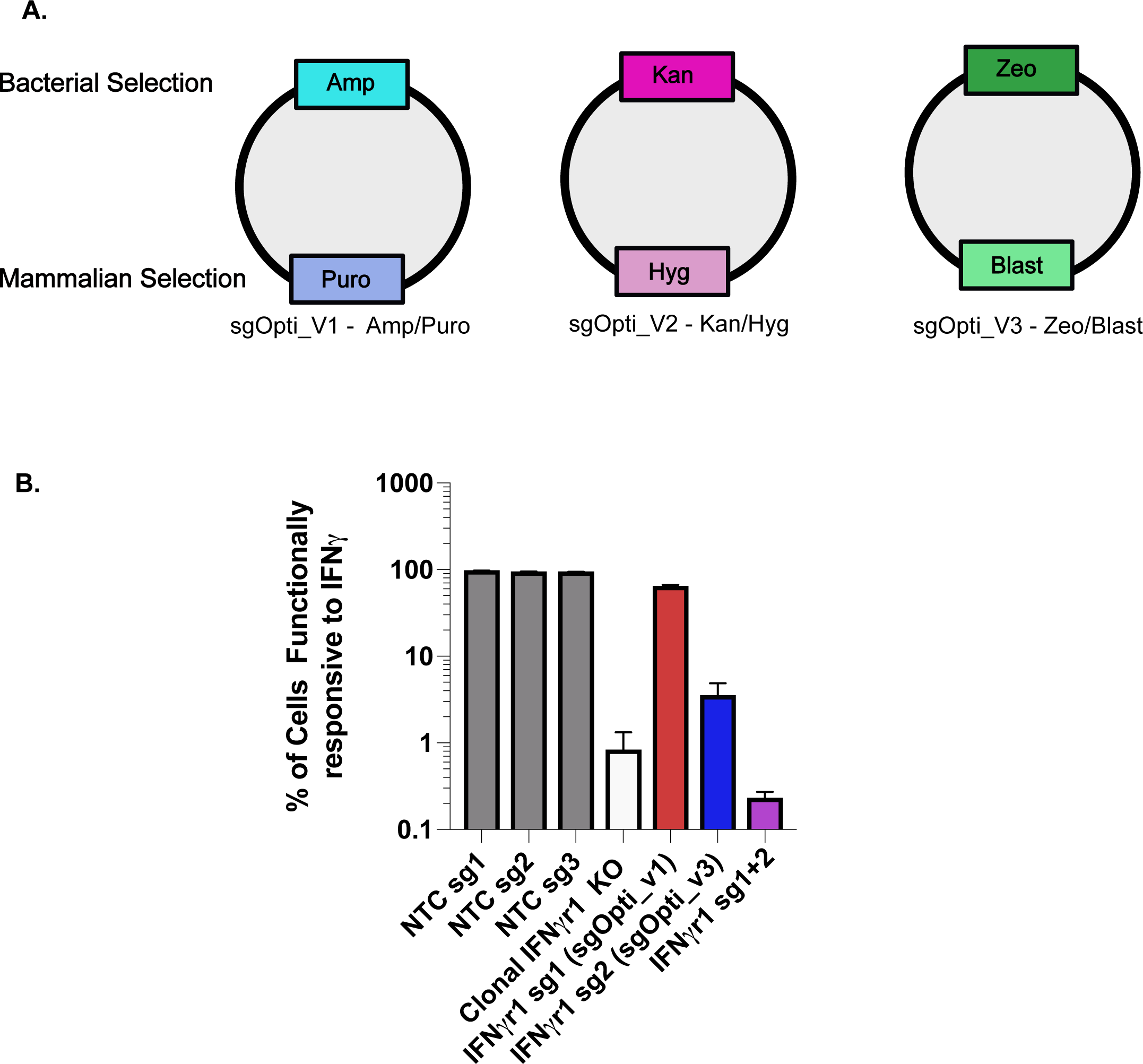
Development of a multi-vector sgRNA system to rapidly edit one gene or simultaneously edit multiple genes. **(A)** Shown is a schematic of the sgOpti derivatives that were generated. sgOpti_V1 was previously published and contains an ampicillin bacterial selection marker and a puromycin mammalian selction marker. sgOpti_V2 and sgOpti_V3 were generated by subcloning distinct bacterial and mammalian selections markers to deliver multiple sgRNAs to cells expressing sgOpti_V1. sgOpti_V2 contains a Kanamycin bacterial selection marker and a Hygromycin B mammalian selection maker while sgOpti_V3 contains a zeocin bacterial selection marker and a Blasticidin mammalian selection marker. **(B)** Cells transduced with the indicated sgRNAs or a clonal IFNγR KO were left untreated or treated with IFNγ for 24 hours and analyzed by flow cytometry for the surface expression of the IFNγ-inducible marker CD271. Shown is the percent of cells that induced CD271 compared to untreated cells for each cell line.

**Figure S5.**
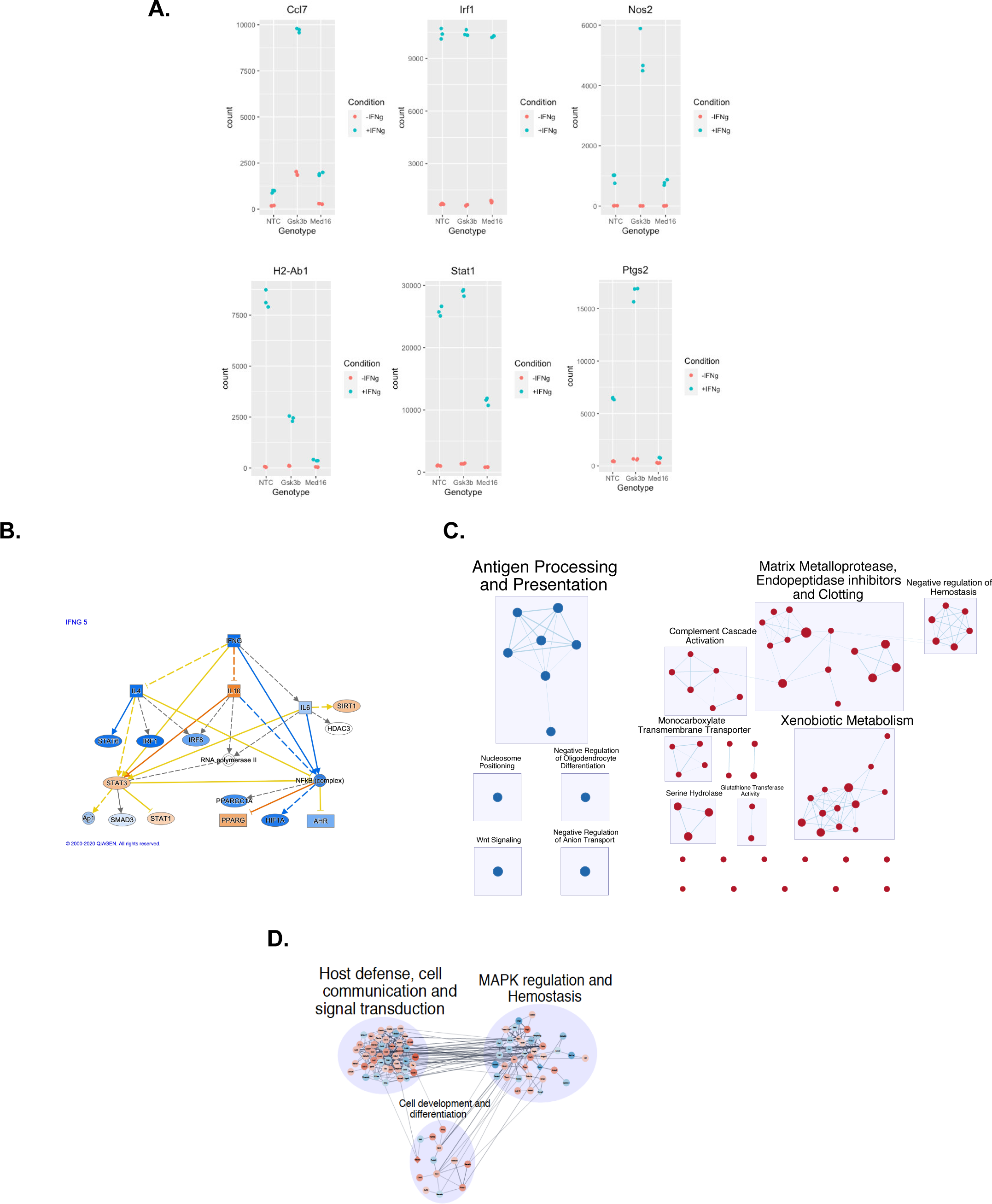
Transcriptomic analysis of MED16 and GSK3β reveals mechanisms of IFNγ-mediated control. **(A)** RNAseq analysis of NTC, Gsk3β KO and Med16 KO cells was completed as described in the materials and methods. Shown are representative scatter plots of normalized absolute read counts for genes that were highly variable among the conditions from the heatmap in Figure 5F. **(B)** Differential gene expression analysis from MED16 KO cells following IFNγ treatment was used to identify dysregulated pathways using gene set enrichment analysis (GSEA). Shown are visual representations of the pathway networks identified using EnrichmentMap and CytoScape. We found a strong downregulation (Blue) of genes involved in antigen processing and presentation and an upregulation (Red) in genes related to Xenobiootic metabolism, glutathione activity, and serine hydrolase and matrix metalloprotease activity. **(C)** GSEA of differentially expressed genes in the Med16 KO after IFNγ-stimulation identified a type I IFN signature. Shown is a pathway map generated by ingenuity pathway analysis highlighting the genes that are downregulated (Blue) or upregulated (Orange) in the Type I IFN pathway. The darkness of the color indicates the magnitude of the differential expression. **(D)** Differential expression analysis of the Gsk3β KO in untreated conditions were used in GSEA. Shown is a visual representation of the dysregulated genes placed into pathway networks using CytoScape. Genes that are upregulated are shown in Red and downregulated genes are shown in Blue. The darkness of the shading indicates the magnitude of the change as determined in the RNAseq analysis.

**Table S1. CRISPR Screen Analysis**

**Table S2. RNAseq Analysis**

**Table S3. Oligonucleotides used in the study**

## Notes

### Competing Interest Statement

The authors have declared no competing interest.

## References

1. van Elsland D, Neefjes J. 2018. Bacterial infections and cancer. EMBO Rep 19.

2. Iwasaki A, Medzhitov R. 2015. Control of adaptive immunity by the innate immune system. Nat Immunol 16:343–53.

3. Tubo NJ, Jenkins MK. 2014. CD4+ T Cells: guardians of the phagosome. Clin Microbiol Rev 27:200–13.

4. DeSandro A, Nagarajan UM, Boss JM. 1999. The bare lymphocyte syndrome: molecular clues to the transcriptional regulation of major histocompatibility complex class II genes. Am J Hum Genet 65:279–86.

5. Reith W, LeibundGut-Landmann S, Waldburger JM. 2005. Regulation of MHC class II gene expression by the class II transactivator. Nat Rev Immunol 5:793–806.

6. Koyama M, Kuns RD, Olver SD, Raffelt NC, Wilson YA, Don AL, Lineburg KE, Cheong M, Robb RJ, Markey KA, Varelias A, Malissen B, Hammerling GJ, Clouston AD, Engwerda CR, Bhat P, MacDonald KP, Hill GR. 2011. Recipient nonhematopoietic antigen-presenting cells are sufficient to induce lethal acute graft-versus-host disease. Nat Med 18:135–42.

7. Abrahimi P, Qin L, Chang WG, Bothwell AL, Tellides G, Saltzman WM, Pober JS. 2016. Blocking MHC class II on human endothelium mitigates acute rejection. JCI Insight 1.

8. Thelemann C, Haller S, Blyszczuk P, Kania G, Rosa M, Eriksson U, Rotman S, Reith W, Acha-Orbea H. 2016. Absence of nonhematopoietic MHC class II expression protects mice from experimental autoimmune myocarditis. Eur J Immunol 46:656–64.

9. Johnson DB, Estrada MV, Salgado R, Sanchez V, Doxie DB, Opalenik SR, Vilgelm AE, Feld E, Johnson AS, Greenplate AR, Sanders ME, Lovly CM, Frederick DT, Kelley MC, Richmond A, Irish JM, Shyr Y, Sullivan RJ, Puzanov I, Sosman JA, Balko JM. 2016. Melanoma-specific MHC-II expression represents a tumour-autonomous phenotype and predicts response to anti-PD-1/PD-L1 therapy. Nat Commun 7:10582.

10. Steimle V, Otten LA, Zufferey M, Mach B. 1993. Complementation cloning of an MHC class II transactivator mutated in hereditary MHC class II deficiency (or bare lymphocyte syndrome). Cell 75:135–46.

11. Jakubzick CV, Randolph GJ, Henson PM. 2017. Monocyte differentiation and antigen-presenting functions. Nat Rev Immunol 17:349–362.

12. Unanue ER, Turk V, Neefjes J. 2016. Variations in MHC Class II Antigen Processing and Presentation in Health and Disease. Annu Rev Immunol 34:265–97.

13. Collins T, Korman AJ, Wake CT, Boss JM, Kappes DJ, Fiers W, Ault KA, Gimbrone MA, Jr., Strominger JL, Pober JS. 1984. Immune interferon activates multiple class II major histocompatibility complex genes and the associated invariant chain gene in human endothelial cells and dermal fibroblasts. Proc Natl Acad Sci U S A 81:4917–21.

14. Neefjes J, Jongsma ML, Paul P, Bakke O. 2011. Towards a systems understanding of MHC class I and MHC class II antigen presentation. Nat Rev Immunol 11:823–36.

15. Buxade M, Huerga Encabo H, Riera-Borrull M, Quintana-Gallardo L, Lopez-Cotarelo P, Tellechea M, Martinez-Martinez S, Redondo JM, Martin-Caballero J, Flores JM, Bosch E, Rodriguez-Fernandez JL, Aramburu J, Lopez-Rodriguez C. 2018. Macrophage-specific MHCII expression is regulated by a remote Ciita enhancer controlled by NFAT5. J Exp Med 215:2901–2918.

16. Ivashkiv LB. 2018. IFNγamma: signalling, epigenetics and roles in immunity, metabolism, disease and cancer immunotherapy. Nat Rev Immunol 18:545–558.

17. Wijdeven RH, van Luijn MM, Wierenga-Wolf AF, Akkermans JJ, van den Elsen PJ, Hintzen RQ, Neefjes J. 2018. Chemical and genetic control of IFNγamma-induced MHCII expression. EMBO Rep 19.

18. Herrero C, Marques L, Lloberas J, Celada A. 2001. IFN-gamma-dependent transcription of MHC class II IA is impaired in macrophages from aged mice. J Clin Invest 107:485–93.

19. Ting JP, Trowsdale J. 2002. Genetic control of MHC class II expression. Cell 109 Suppl:S21–33.

20. Bousoik E, Montazeri Aliabadi H. 2018. “Do We Know Jack” About JAK? A Closer Look at JAK/STAT Signaling Pathway. Front Oncol 8:287.

21. Hu X, Ivashkiv LB. 2009. Cross-regulation of signaling pathways by interferon-gamma: implications for immune responses and autoimmune diseases. Immunity 31:539–50.

22. Schroder K, Hertzog PJ, Ravasi T, Hume DA. 2004. Interferon-gamma: an overview of signals, mechanisms and functions. J Leukoc Biol 75:163–89.

23. Lehtonen A, Matikainen S, Julkunen I. 1997. Interferons up-regulate STAT1, STAT2, and IRF family transcription factor gene expression in human peripheral blood mononuclear cells and macrophages. J Immunol 159:794–803.

24. Beresford GW, Boss JM. 2001. CIITA coordinates multiple histone acetylation modifications at the HLA-DRA promoter. Nat Immunol 2:652–7.

25. Paul P, van den Hoorn T, Jongsma ML, Bakker MJ, Hengeveld R, Janssen L, Cresswell P, Egan DA, van Ham M, Ten Brinke A, Ovaa H, Beijersbergen RL, Kuijl C, Neefjes J. 2011. A Genome-wide multidimensional RNAi screen reveals pathways controlling MHC class II antigen presentation. Cell 145:268–83.

26. Oh J, Wu N, Baravalle G, Cohn B, Ma J, Lo B, Mellman I, Ishido S, Anderson M, Shin JS. 2013. MARCH1-mediated MHCII ubiquitination promotes dendritic cell selection of natural regulatory T cells. J Exp Med 210:1069–77.

27. Alix E, Godlee C, Cerny O, Blundell S, Tocci R, Matthews S, Liu M, Pruneda JN, Swatek KN, Komander D, Sleap T, Holden DW. 2020. The Tumour Suppressor TMEM127 Is a Nedd4-Family E3 Ligase Adaptor Required by Salmonella SteD to Ubiquitinate and Degrade MHC Class II Molecules. Cell Host Microbe 28:54–68 e7.

28. Ankley L, Thomas S, Olive AJ. 2020. Fighting Persistence: How Chronic Infections with Mycobacterium tuberculosis Evade T Cell-Mediated Clearance and New Strategies To Defeat Them. Infect Immun 88.

29. Grau V, Herbst B, Steiniger B. 1998. Dynamics of monocytes/macrophages and T lymphocytes in acutely rejecting rat renal allografts. Cell Tissue Res 291:117–26.

30. Underhill DM, Bassetti M, Rudensky A, Aderem A. 1999. Dynamic interactions of macrophages with T cells during antigen presentation. J Exp Med 190:1909–14.

31. Pai RK, Convery M, Hamilton TA, Boom WH, Harding CV. 2003. Inhibition of IFN-gamma-induced class II transactivator expression by a 19-kDa lipoprotein from Mycobacterium tuberculosis: a potential mechanism for immune evasion. J Immunol 171:175–84.

32. Pennini ME, Pai RK, Schultz DC, Boom WH, Harding CV. 2006. Mycobacterium tuberculosis 19-kDa lipoprotein inhibits IFN-gamma-induced chromatin remodeling of MHC2TA by TLR2 and MAPK signaling. J Immunol 176:4323–30.

33. Zhong G, Fan T, Liu L. 1999. Chlamydia inhibits interferon gamma-inducible major histocompatibility complex class II expression by degradation of upstream stimulatory factor 1. J Exp Med 189:1931–8.

34. Brinkman EK, Chen T, Amendola M, van Steensel B. 2014. Easy quantitative assessment of genome editing by sequence trace decomposition. Nucleic Acids Res 42:e168.

35. Steimle V, Durand B, Barras E, Zufferey M, Hadam MR, Mach B, Reith W. 1995. A novel DNA-binding regulatory factor is mutated in primary MHC class II deficiency (bare lymphocyte syndrome). Genes Dev 9:1021–32.

36. Doench JG, Fusi N, Sullender M, Hegde M, Vaimberg EW, Donovan KF, Smith I, Tothova Z, Wilen C, Orchard R, Virgin HW, Listgarten J, Root DE. 2016. Optimized sgRNA design to maximize activity and minimize off-target effects of CRISPR-Cas9. Nat Biotechnol 34:184–191.

37. Hart T, Brown KR, Sircoulomb F, Rottapel R, Moffat J. 2014. Measuring error rates in genomic perturbation screens: gold standards for human functional genomics. Mol Syst Biol 10:733.

38. Li W, Xu H, Xiao T, Cong L, Love MI, Zhang F, Irizarry RA, Liu JS, Brown M, Liu XS. 2014. MAGeCK enables robust identification of essential genes from genome-scale CRISPR/Cas9 knockout screens. Genome Biol 15:554.

39. Kanehisa M, Goto S. 2000. KEGG: kyoto encyclopedia of genes and genomes. Nucleic Acids Res 28:27–30.

40. Kanehisa M, Sato Y, Furumichi M, Morishima K, Tanabe M. 2019. New approach for understanding genome variations in KEGG. Nucleic Acids Res 47:D590–D595.

41. Kanehisa M. 2019. Toward understanding the origin and evolution of cellular organisms. Protein Sci 28:1947–1951.

42. Poss ZC, Ebmeier CC, Taatjes DJ. 2013. The Mediator complex and transcription regulation. Crit Rev Biochem Mol Biol 48:575–608.

43. Wu D, Pan W. 2010. GSK3: a multifaceted kinase in Wnt signaling. Trends Biochem Sci 35:161–8.

44. Conaway RC, Conaway JW. 2011. Origins and activity of the Mediator complex. Semin Cell Dev Biol 22:729–34.

45. Beurel E, Michalek SM, Jope RS. 2010. Innate and adaptive immune responses regulated by glycogen synthase kinase-3 (GSK3). Trends Immunol 31:24–31.

46. Thomson AW, Turnquist HR, Raimondi G. 2009. Immunoregulatory functions of mTOR inhibition. Nat Rev Immunol 9:324–37.

47. Xu Y, Harton JA, Smith BD. 2008. CIITA mediates interferon-gamma repression of collagen transcription through phosphorylation-dependent interactions with co-repressor molecules. J Biol Chem 283:1243–56.

48. An WF, Germain AR, Bishop JA, Nag PP, Metkar S, Ketterman J, Walk M, Weiwer M, Liu X, Patnaik D, Zhang YL, Gale J, Zhao W, Kaya T, Barker D, Wagner FF, Holson EB, Dandapani S, Perez J, Munoz B, Palmer M, Pan JQ, Haggarty SJ, Schreiber SL. 2010. Discovery of Potent and Highly Selective Inhibitors of GSK3β, Probe Reports from the NIH Molecular Libraries Program, Bethesda (MD).

49. Ring DB, Johnson KW, Henriksen EJ, Nuss JM, Goff D, Kinnick TR, Ma ST, Reeder JW, Samuels I, Slabiak T, Wagman AS, Hammond ME, Harrison SD. 2003. Selective glycogen synthase kinase 3 inhibitors potentiate insulin activation of glucose transport and utilization in vitro and in vivo. Diabetes 52:588–95.

50. Wang GG, Calvo KR, Pasillas MP, Sykes DB, Hacker H, Kamps MP. 2006. Quantitative production of macrophages or neutrophils ex vivo using conditional Hoxb8. Nat Methods 3:287–93.

51. Beurel E, Jope RS. 2008. Differential regulation of STAT family members by glycogen synthase kinase-3. J Biol Chem 283:21934–44.

52. Huang J, Guo X, Li W, Zhang H. 2017. Activation of Wnt/beta-catenin signalling via GSK3 inhibitors direct differentiation of human adipose stem cells into functional hepatocytes. Sci Rep 7:40716.

53. Fulco CP, Munschauer M, Anyoha R, Munson G, Grossman SR, Perez EM, Kane M, Cleary B, Lander ES, Engreitz JM. 2016. Systematic mapping of functional enhancer-promoter connections with CRISPR interference. Science 354:769–773.

54. Subramanian A, Tamayo P, Mootha VK, Mukherjee S, Ebert BL, Gillette MA, Paulovich A, Pomeroy SL, Golub TR, Lander ES, Mesirov JP. 2005. Gene set enrichment analysis: a knowledge-based approach for interpreting genome-wide expression profiles. Proc Natl Acad Sci U S A 102:15545–50.

55. Roan NR, Gierahn TM, Higgins DE, Starnbach MN. 2006. Monitoring the T cell response to genital tract infection. Proc Natl Acad Sci U S A 103:12069–74.

56. Siska PJ, Rathmell JC. 2016. Metabolic Signaling Drives IFN-gamma. Cell Metab 24:651–652.

57. Garg S, Sharma M, Ung C, Tuli A, Barral DC, Hava DL, Veerapen N, Besra GS, Hacohen N, Brenner MB. 2011. Lysosomal trafficking, antigen presentation, and microbial killing are controlled by the Arf-like GTPase Arl8b. Immunity 35:182–93.

58. Michelet X, Garg S, Wolf BJ, Tuli A, Ricciardi-Castagnoli P, Brenner MB. 2015. MHC class II presentation is controlled by the lysosomal small GTPase, Arl8b. J Immunol 194:2079–88.

59. Sekine H, Okazaki K, Ota N, Shima H, Katoh Y, Suzuki N, Igarashi K, Ito M, Motohashi H, Yamamoto M. 2016. The Mediator Subunit MED16 Transduces NRF2-Activating Signals into Antioxidant Gene Expression. Mol Cell Biol 36:407–20.

60. Bancerek J, Poss ZC, Steinparzer I, Sedlyarov V, Pfaffenwimmer T, Mikulic I, Dolken L, Strobl B, Muller M, Taatjes DJ, Kovarik P. 2013. CDK8 kinase phosphorylates transcription factor STAT1 to selectively regulate the interferon response. Immunity 38:250–62.

61. Smith JL, Jeng S, McWeeney SK, Hirsch AJ. 2017. A MicroRNA Screen Identifies the Wnt Signaling Pathway as a Regulator of the Interferon Response during Flavivirus Infection. J Virol 91.

62. Bai M, Li W, Yu N, Zhang H, Long F, Zeng A. 2017. The crosstalk between beta-catenin signaling and type I, type II and type III interferons in lung cancer cells. Am J Transl Res 9:2788–2797.

63. Huang WC, Lin YS, Wang CY, Tsai CC, Tseng HC, Chen CL, Lu PJ, Chen PS, Qian L, Hong JS, Lin CF. 2009. Glycogen synthase kinase-3 negatively regulates anti-inflammatory interleukin-10 for lipopolysaccharide-induced iNOS/NO biosynthesis and RANTES production in microglial cells. Immunology 128:e275–86.

64. Garvin AJ, Khalaf AHA, Rettino A, Xicluna J, Butler L, Morris JR, Heery DM, Clarke NM. 2019. GSK3βeta-SCFFBXW7alpha mediated phosphorylation and ubiquitination of IRF1 are required for its transcription-dependent turnover. Nucleic Acids Res 47:4476–4494.

65. Gibbs KD, Washington EJ, Jaslow SL, Bourgeois JS, Foster MW, Guo R, Brennan RG, Ko DC. 2020. The Salmonella Secreted Effector SarA/SteE Mimics Cytokine Receptor Signaling to Activate STAT3. Cell Host Microbe 27:129–139 e4.

66. Panagi I, Jennings E, Zeng J, Gunster RA, Stones CD, Mak H, Jin E, Stapels DAC, Subari NZ, Pham THM, Brewer SM, Ong SYQ, Monack DM, Helaine S, Thurston TLM. 2020. Salmonella Effector SteE Converts the Mammalian Serine/Threonine Kinase GSK3 into a Tyrosine Kinase to Direct Macrophage Polarization. Cell Host Microbe 27:41–53 e6.

67. Garcia-Diaz A, Shin DS, Moreno BH, Saco J, Escuin-Ordinas H, Rodriguez GA, Zaretsky JM, Sun L, Hugo W, Wang X, Parisi G, Saus CP, Torrejon DY, Graeber TG, Comin-Anduix B, Hu-Lieskovan S, Damoiseaux R, Lo RS, Ribas A. 2017. Interferon Receptor Signaling Pathways Regulating PD-L1 and PD-L2 Expression. Cell Rep 19:1189–1201.

68. Mandai M, Hamanishi J, Abiko K, Matsumura N, Baba T, Konishi I. 2016. Dual Faces of IFNγamma in Cancer Progression: A Role of PD-L1 Induction in the Determination of Pro- and Antitumor Immunity. Clin Cancer Res 22:2329–34.

69. Gu W, Chen J, Yang L, Zhao KN. 2012. TNF-alpha promotes IFN-gamma-induced CD40 expression and antigen process in Myb-transformed hematological cells. ScientificWorldJournal 2012:621969.

70. Zhou F. 2009. Molecular mechanisms of IFN-gamma to up-regulate MHC class I antigen processing and presentation. Int Rev Immunol 28:239–60.

71. Shalem O, Sanjana NE, Hartenian E, Shi X, Scott DA, Mikkelson T, Heckl D, Ebert BL, Root DE, Doench JG, Zhang F. 2014. Genome-scale CRISPR-Cas9 knockout screening in human cells. Science 343:84–87.

72. Reimand J, Isserlin R, Voisin V, Kucera M, Tannus-Lopes C, Rostamianfar A, Wadi L, Meyer M, Wong J, Xu C, Merico D, Bader GD. 2019. Pathway enrichment analysis and visualization of omics data using g:Profiler, GSEA, Cytoscape and EnrichmentMap. Nat Protoc 14:482–517.

